# A chemical-genetic map of the pathways controlling drug potency in *Mycobacterium tuberculosis*

**DOI:** 10.1101/2021.11.27.469863

**Authors:** Shuqi Li, Nicholas C. Poulton, Jesseon S. Chang, Zachary A. Azadian, Michael A. DeJesus, Nadine Ruecker, Matthew D. Zimmerman, Kathryn Eckartt, Barbara Bosch, Curtis Engelhart, Daniel Sullivan, Martin Gengenbacher, Véronique A. Dartois, Dirk Schnappinger, Jeremy M. Rock

## Abstract

*Mycobacterium tuberculosis* (Mtb) infection is notoriously difficult to treat. Treatment efficacy is limited by Mtb’s intrinsic drug resistance, as well as its ability to evolve acquired resistance to all antituberculars in clinical use. A deeper understanding of the bacterial pathways that govern drug efficacy could facilitate the development of more effective therapies to overcome resistance, identify new mechanisms of acquired resistance, and reveal overlooked therapeutic opportunities. To define these pathways, we developed a CRISPR interference chemical-genetics platform to titrate the expression of Mtb genes and quantify bacterial fitness in the presence of different drugs. Mining this dataset, we discovered diverse and novel mechanisms of intrinsic drug resistance, unveiling hundreds of potential targets for synergistic drug combinations. Combining chemical-genetics with comparative genomics of Mtb clinical isolates, we further identified numerous new potential mechanisms of acquired drug resistance, one of which is associated with the emergence of a multidrug-resistant tuberculosis (TB) outbreak in South America. Lastly, we make the unexpected discovery of an “acquired drug sensitivity.” We found that the intrinsic resistance factor *whiB7* was inactivated in an entire Mtb sublineage endemic to Southeast Asia, presenting an opportunity to potentially repurpose the macrolide antibiotic clarithromycin to treat TB. This chemical-genetic map provides a rich resource to understand drug efficacy in Mtb and guide future TB drug development and treatment.

## INTRODUCTION

Infections caused by the bacterial pathogen *Mycobacterium tuberculosis* (Mtb) are notoriously difficult to treat. Current standard of care requires a multidrug regimen lasting for several months, which limits patient compliance and contributes to the development of drug-resistant Mtb (WHO, 2021). While the reasons necessitating prolonged chemotherapy are multifactorial, including variable drug penetration into Mtb-containing granulomas (Dartois, 2014) and the presence of phenotypically drug-tolerant bacterial subpopulations (Balaban et al., 2019), the intrinsic resistance of the infecting bacterium and its ability to evolve acquired resistance to all antituberculars in clinical use limits treatment efficacy (Colangeli et al., 2018; Xu et al., 2017).

Mtb is intrinsically resistant to many antibacterials. While relatively underexplored, intrinsic resistance is typically ascribed to the low permeability of the Mtb cell envelope and the numerous efflux pumps encoded in the Mtb genome (Batt et al., 2020; Jarlier and Nikaido, 1994; da Silva et al., 2011). All acquired drug resistance in Mtb occurs via mutation, and in recent decades many resistance mutations have been mapped and characterized (Walker et al., 2015). These mutations most commonly occur in the drug target or drug activator, reducing the affinity of the drug-target interaction or reducing conversion of the drug to the bioactive molecule (Banerjee et al., 1994; Walker et al., 2015; Zhang et al., 1992). Yet, our knowledge of acquired drug resistance in Mtb remains incomplete, particularly for mutations outside of the drug target or activator and which typically confer low to intermediate, but clinically relevant, levels of drug resistance (Carter, 2021; Colangeli et al., 2018; Hicks et al., 2020; Walker et al., 2015). A deeper understanding of both intrinsic and acquired drug resistance in Mtb could facilitate the development of therapies to overcome resistance mechanisms, improve the diagnosis of drug-resistant TB, and reveal overlooked therapeutic opportunities (Blondiaux et al., 2017; Hugonnet et al., 2009).

To provide a genome-wide overview of the bacterial pathways that control drug potency, we developed a CRISPR interference (CRISPRi) (Bosch et al., 2021; Choudhary et al., 2015; Qi et al., 2013; Rock et al., 2017) chemical-genetics platform to titrate the expression of nearly all Mtb genes (essential and non-essential) and quantify bacterial fitness in the presence of different drugs. This approach identified hundreds of Mtb genes whose inhibition altered fitness in the presence of partially inhibitory drug concentrations, including genes encoding the direct drug target and non-target hit genes. Mining this dataset, we discovered diverse mechanisms of intrinsic drug resistance that can be targeted to potentiate therapy. Overlaying the chemical-genetic results with comparative genomics of Mtb clinical isolates, we identified new, clinically relevant mechanisms of acquired drug resistance. Lastly, we make the unexpected discovery of “acquired drug sensitivities,” whereby loss-of-function mutations in intrinsic drug resistance genes render some Mtb clinical strains hypersusceptible to clarithromycin, a macrolide antibiotic not typically used to treat tuberculosis (TB). This chemical-genetic map provides a rich resource to understand drug potency in Mtb and guide future TB drug development and treatment.

## RESULTS

To define genes that alter drug potency in Mtb, we performed 90 CRISPRi screens across nine drugs in the Mtb reference strain, H37Rv. These screens used a genome-scale CRISPRi library containing 96,700 sgRNAs (Bosch et al., 2021) to enable titratable knockdown for nearly all Mtb genes, including both protein coding genes and non-coding RNAs (**Figure 1A**). Titrated gene knockdown was achieved by targeting non-canonical Sth1 dCas9 protospacer adjacent motifs (PAMs) (Rock et al., 2017) and modulating the extent of complementarity between the sgRNA and DNA target (Bosch et al., 2021). Knockdown tuning enabled hypomorphic silencing of *in vitro* essential genes, thereby allowing assessment of chemical-genetic interactions for both *in vitro* essential and non-essential genes to provide a global overview of gene-drug interactions in Mtb.

**Figure 1:**
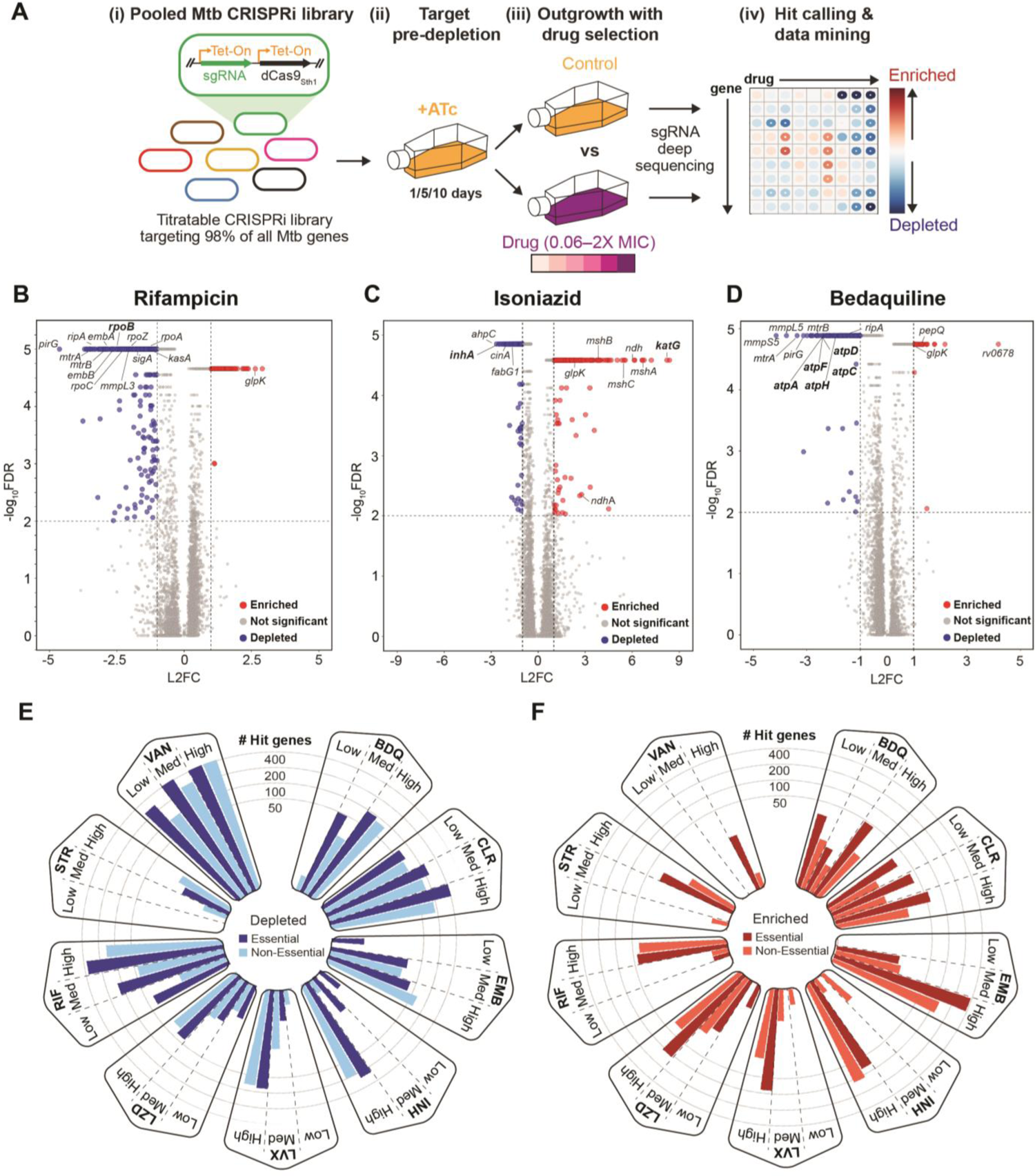
Chemical-genetic profiling identifies hundreds of genes that alter drug efficacy in *M. tuberculosis*. (A) Experimental design to quantify chemical-genetic interactions in Mtb. (i) The pooled Mtb CRISPRi library contains 96,700 sgRNAs targeting 4,052/4,125 of all Mtb genes. *In vitro* essential genes were targeted for titratable knockdown by varying the targeted PAM and sgRNA targeting sequence length; non-essential genes were targeted only with the strongest available sgRNAs (Bosch et al., 2021). (ii) The CRISPRi inducer anhydrotetracycline (ATc) was added for 1, 5, or 10 days prior to drug exposure to pre-deplete target gene products. (iii) Triplicate cultures were outgrown +ATc in DMSO or drug at six concentrations spanning the predicted minimum inhibitory concentration (MIC). (iv) Following outgrowth, genomic DNA was harvested from cultures treated with three descending doses of partially inhibitory drug concentrations (“High”, “Med”, and “Low”; **Supplemental Figure 1**), sgRNA targeting sequences amplified for next-generation sequencing, and hit genes called with MAGeCK. (B-D) Volcano plots showing log2 fold-change (L2FC) values and false discovery rates (FDR) for each gene after culture outgrowth in the presence of the indicated drugs. Results for the highest partially inhibitory concentration (“High”; **Supplemental Figure 1**) for the 5-day CRISPRi library pre-depletion screen are shown. (E-F) The number of significantly depleted and enriched hit genes (FDR < 0.01, |L2FC| > 1; union of 1 and 5-day CRISPRi library pre-depletion screens) are shown for the indicated drugs and concentrations (**Supplemental Figure 1**). Gene essentiality calls were defined by CRISPRi as in (Bosch et al., 2021).

Anhydrotetracycline (ATc) was added 1, 5, or 10 days prior to drug exposure to transcriptionally activate CRISPRi and deplete target gene products (**Figure 1A**). Drugs were chosen to represent the majority of clinically relevant Mtb targets (**Table 1**), including three of the four first-line agents (pyrazinamide was not included because it is not active under standard culture conditions), four second-line agents, and two drugs not traditionally used to treat TB. Drugs were screened at concentrations spanning the predicted minimum inhibitory concentration (MIC) (**Supplemental Figure 1A-I**). Triplicate CRISPRi library cultures were outgrown in the presence or absence of drug. After outgrowth, we harvested genomic DNA from cultures treated with three descending doses of partially inhibitory drug concentrations (“High”, “Med”, and “Low”; **Supplemental Figure 1A-I**) and analyzed sgRNA abundance by deep sequencing. Growth phenotypes were well correlated among triplicate screens (average Pearson correlation between replicate screens: r > 0.99). Hits were identified by MAGeCK (Li et al., 2014) as those genes whose CRISPRi inhibition reduced or increased relative fitness in the presence of a given drug (false discovery rate (FDR) < 0.01, log2 fold-change |L2FC| > 1). Analysis of the number of unique hit genes across different drugs showed that the 1 and 5-day target pre-depletion datasets recovered the majority (>95%) of unique hits (**Supplemental Figure 2A-I; Supplemental Data 1**). Thus, hit genes were further defined as the union of 1 and 5-day target pre-depletion screens. These criteria identified 1,373 genes whose knockdown led to sensitization and 775 genes whose knockdown led to resistance to at least one drug (**Supplemental Data 1**), representing at least one chemical-genetic interaction for 38.5% (n=1,587/4,125) of all annotated genes in the Mtb genome. Most hit genes had a single chemical-genetic interaction, but some had as many as seven (**Supplemental Figure 2J, Supplemental Data 1**).

**Table 1:**
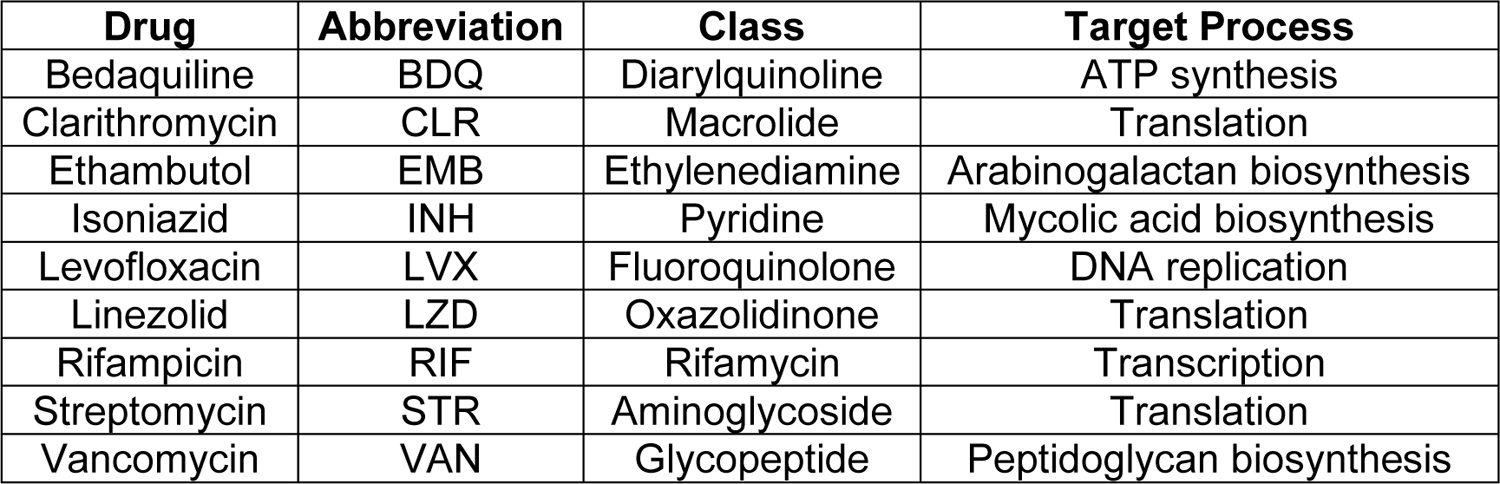
Drugs Profiled and their Target Processes

| Drug | Abbreviation | Class | Target Process |
| --- | --- | --- | --- |
| Bedaquiline | BDQ | Diarylquinoline | ATP synthesis |
| Clarithromycin | CLR | Macrolide | Translation |
| Ethambutol | EMB | Ethylenediamine | Arabinogalactan biosynthesis |
| Isoniazid | INH | Pyridine | Mycolic acid biosynthesis |
| Levofloxacin | LVX | Fluoroquinolone | DNA replication |
| Linezolid | LZD | Oxazolidinone | Translation |
| Rifampicin | RIF | Rifamycin | Transcription |
| Streptomycin | STR | Aminoglycoside | Translation |
| Vancomycin | VAN | Glycopeptide | Peptidoglycan biosynthesis |

The chemical-genetic screens recovered expected hit genes. For example, the genes encoding the targets of rifampicin (*rpoB*) (Campbell et al., 2001) (**Figure 1B**), isoniazid (*inhA*) (Banerjee et al., 1994) (**Figure 1C**), and bedaquiline (ATP synthase) (Andries et al., 2005) (**Figure 1D**) were among the most sensitized hits in each respective screen. Genes encoding the targets of known synergistic drug combinations were also recovered, for example ethambutol (*embAB*) + rifampicin and SQ109 (*mmpL3*) + rifampicin (**Figure 1B**) (Cokol et al., 2017). Lastly, genes whose inactivation is known to confer acquired drug resistance were also observed, including glycerol kinase *glpK* (**Figure 1B-D**) (Bellerose et al., 2019; Safi et al., 2019), catalase-peroxidase *katG* (**Figure 1C**) (Vilchèze and Jacobs JR., 2014), and the transcriptional repressor *rv0678* (**Figure 1D**) (Andries et al., 2014). Consistent with the robust recovery of expected hits, benchmarking our CRISPRi approach against published transposon sequencing (TnSeq) chemical-genetic results revealed a high degree of overlap (63.3-87.7% TnSeq hit recovery; **Supplemental Data 2**) (Xu et al., 2017), although TnSeq is necessarily restricted to interrogation of *in vitro* non-essential genes, at least as currently implemented in Mtb.

The number of hit genes varied widely across drugs (**Figure 1E,F**; **Supplemental Data 1; Supplemental Figure 2A-I;**), from hundreds for rifampicin, vancomycin, and ethambutol to tens for streptomycin. Interestingly, *in vitro* essential genes were enriched relative to non-essential genes for chemical-genetic interactions (**Figure 1E,F; Supplemental Figure 3A**), even when taking into account the bias towards sgRNAs targeting *in vitro* essential genes in the CRISPRi library. This enrichment demonstrates the increased information content available when assaying essential genes by chemical-genetics and highlights the power of titratable CRISPRi to assay gene classes typically intractable with more traditional approaches like TnSeq. Hierarchical clustering of hit genes revealed unique chemical-genetic signatures for each drug (**Supplemental Figure 3B**) that were then mined for biological insight.

Despite the fact that they target distinct cellular processes (**Table 1**), clustering analysis revealed correlated chemical-genetic signatures for rifampicin, vancomycin, and bedaquiline (**Supplemental Figure 3B**), suggesting shared mechanisms of intrinsic resistance or sensitivity. Enrichment analysis identified the essential mycolic acid-arabinogalactan-peptidoglycan (mAGP) complex to be a common sensitizing hit between rifampicin, vancomycin, and bedaquiline but not the ribosome targeting drugs clarithromycin, linezolid, or streptomycin (**Supplemental Figure 3C**). The mAGP is the primary constituent of the cell envelope and has long been known to serve as a permeability barrier that mediates intrinsic drug resistance in Mtb (Batt et al., 2020; Jarlier and Nikaido, 1994; da Silva et al., 2011). Interestingly, it was not obvious which chemical features for each drug were driving selective sensitization to mAGP disruption (**Supplemental Figure 3D**) (Davis et al., 2014). For example, despite having similar molecular weights, bedaquiline (555.5 daltons) displayed a strong mAGP signature whereas streptomycin (581.6 daltons) did not; despite similar polar surface areas, rifampicin (220 Å^2^) displays a strong mAGP signature but clarithromycin (183 Å^2^) does not.

To ensure the validity of the screen results, we quantified drug susceptibility with individual hypomorphic CRISPRi strains targeting mAGP-biosynthetic genes, demonstrating 2- to 43-fold reductions in IC_50_ for rifampicin, vancomycin, and bedaquiline but little to no change in IC_50_ for linezolid (**Supplemental Figure 4A-D**). To validate these results chemically, we chose to focus on the β-ketoacyl-ACP synthase KasA. KasA is an essential component of the FAS-II pathway responsible for elongation of meromycolic acids and is an actively pursued drug target (Abrahams et al., 2016; Kumar et al., 2018). Consistent with our genetic results, checkerboard assays demonstrated synergy between the KasA inhibitor GSK3011724A (GSK’724A) and rifampicin, vancomycin, and bedaquiline but not linezolid (**Supplemental Figure 4E,F**).

Because drug interactions can be influenced by the growth environment (Lenaerts et al., 2015), we also confirmed the synergy between GSK’724A and rifampicin in a macrophage infection model (**Supplemental Figure 4G**). Consistent with the hypothesis that synergy between GSK’724A and this mechanistically diverse group of antibiotics could be explained, at least in part, by inhibition of mycolic acid biosynthesis resulting in improved drug uptake, Mtb cultures pre-treated with a sub-MIC dose of GSK’724A showed increased uptake of ethidium bromide and a fluorescent vancomycin conjugate (**Supplemental Figure 4H,I**). These results validate the screen and confirm the role of the mAGP complex as a selective mechanism of intrinsic resistance relevant for some antitubercular agents but not others (Larrouy-Maumus et al., 2016).

### The two-component system *mtrAB* and associated lipoprotein *lpqB* promote envelope integrity and are central mediators of intrinsic drug resistance in Mtb

Two of the most sensitizing hit genes across multiple drugs were the response regulator *mtrA* and its cognate histidine kinase *mtrB* (**Figure 1B,D; Figure 2A; Supplemental Data 1**), which together encode the MtrAB two-component signaling system (Gorla et al., 2018; Zahrt and Deretic, 2000). The *mtrAB* operon also encodes a putative lipoprotein *lpqB* (**Figure 2B**). LpqB is proposed to interact with MtrB to promote MtrA phosphorylation and activation (Nguyen et al., 2010). The similarities between the chemical-genetic signatures of *mtrAB-lpqB* and mAGP biosynthetic genes (**Figure 2A; Supplemental Figure 4B**) are consistent with a critical role for this two-component system in regulating mAGP integrity. Given the predicted essentiality of *mtrA*, *mtrB*, and *lpqB* (Dejesus et al., 2017; Zahrt and Deretic, 2000) and the magnitude by which inhibition of these genes sensitized Mtb to various antibiotics, we next sought to better define the mechanism by which *mtrAB-lpqB* promotes intrinsic drug resistance.

**Figure 2:**
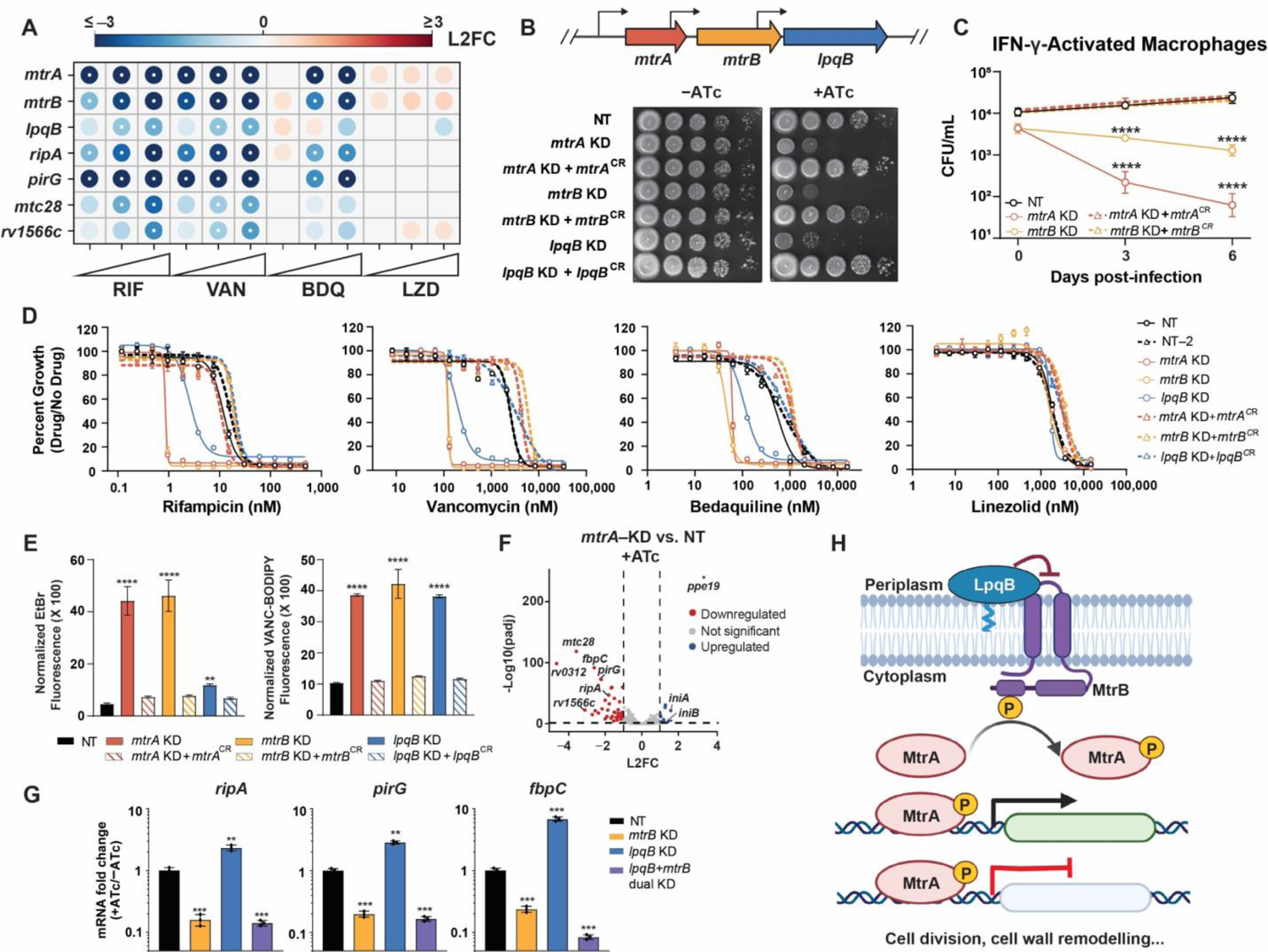
MtrAB-LpqB promote envelope integrity and are central mediators of intrinsic drug resistance. (A) Feature-expression heatmap of select chemical-genetic hit genes for the indicated drugs from the 5-day CRISPRi library pre-depletion screen. The color of each circle represents the gene-level L2FC. A white dot represents an FDR < 0.01 and a |L2FC| > 1. (B) Growth of the indicated CRISPRi strains. NT = non-targeting; KD = knockdown; CR = CRISPRi-resistant. Transcriptional start sites (Shell et al., 2015) are indicated with black arrows. (C) Growth of the indicated CRISPRi strains in IFN-γ-activated murine bone marrow derived macrophages. Bacterial strains were exposed to ATc for 24 hours prior to macrophage infection. 3 and 6 days after infection, bacteria were harvested and quantified by colony-forming units (CFU). Data represent mean ± SEM for technical triplicates. Significance was determined by two-way ANOVA and adjusted for multiple comparisons. ****, p<0.0001. (D) MIC values for the indicated drugs were measured against the indicated strains. Data represent mean ± SEM for technical triplicates and are representative of at least two independent experiments. (E) Ethidium bromide and Vancomycin-BODIPY uptake of the indicated strains. Data represent mean ± SEM for three replicates and are representative of at least two independent experiments. Results from an unpaired t-test are shown: ****, p<0.0001. (F) *mtrA* and NT CRISPRi strains were grown for two days with ATc, after which RNA was harvested and sequenced. L2FC and –log10 (p_adj_) for each gene are plotted. Dashed lines mark significant hits (p_adj_ < 0.05 and |L2FC| > 1). (G) Quantification of indicated gene mRNA levels by qRT-PCR. Strains were grown in the presence or absence of ATc for ∼3 generations prior to harvesting RNA. Error bars are SEM of three technical replicates. Results from an unpaired t-test are shown: **, p <0.01; ***, p<0.001. (H) Schematic of the proposed MtrAB-LpqB signaling system. The histidine kinase MtrB activates the response regulator MtrA to control expression of genes important for proper cell division and cell wall remodeling. The lipoprotein LpqB interacts with MtrB and may negatively regulate MtrB-dependent activation of MtrA.

Consistent with the predicted essentiality, strong CRISPRi-silencing of *mtrA*, *mtrB*, and *lpqB* prevented Mtb growth (**Figure 2B**). Complementation of CRISPRi knockdown with CRISPRi-resistant *mtrA*, *mtrB*, or *lpqB* alleles reversed this growth defect (**Figure 2B**), demonstrating specificity for the observed phenotypes.

Whereas inhibition of *mtrB* was bacteriostatic both in axenic culture and macrophages, inhibition of *mtrA* was bacteriostatic in axenic culture but bactericidal in resting and IFN-γ activated macrophages (**Figure 2C, Supplemental Figure 5A,B**). Consistent with the chemical-genetic screen results, knockdown of *mtrA*, *mtrB*, and *lpqB* strongly sensitized Mtb (10-100 fold decreases in IC_50_) to rifampicin, vancomycin, and bedaquiline, but not other drugs (**Figure 2D, Supplemental Figure 5C**). Also consistent with the screen, the magnitude of drug sensitization was not identical across all three genes, being more similar between *mtrA* and *mtrB* than *lpqB* (**Figure 2D**). As with inhibition of KasA (**Supplemental Figure 4H,I**), silencing of *mtrA*, *mtrB*, and to a lesser extent *lpqB* led to increased permeability to ethidium bromide and a fluorescent vancomycin conjugate (**Figure 2E**). Together, these results suggest that the increase in drug susceptibility in *mtrAB-lpqB* knockdown strains is at least in part mediated by increased envelope permeability and are consistent with an essential role for *mtrAB-lpqB* in regulating mAGP integrity.

To better understand the mechanism(s) by which the response regulator *mtrA* mediates multi-drug intrinsic resistance, we next defined its regulon. RNA-sequencing (RNA-seq) following *mtrA* silencing identified 41 significantly down-regulated and 11 significantly upregulated genes (p_adi_ < 0.05, |L2FC| > 1) (**Figure 2F**, **Supplemental Data 3**). Consistent with a direct regulatory role, MtrA was found to bind the promoters of 25 of the significantly down-regulated and two of the significantly upregulated genes in a previously published ChIP-seq study (**Supplemental Figure 5D**, **Supplemental Data 3**) (Gorla et al., 2018). Upregulated genes not found to be bound by MtrA included the envelope stress response genes *iniB* and *iniA* (Alland et al., 2000), and thus at least some of the upregulated genes following *mtrA* silencing may reflect secondary consequences of envelope stress. A consensus MtrA recognition site derived from promoters of genes differentially regulated upon *mtrA* inhibition and found to interact with MtrA by ChIP-seq was broadly similar to previously published MtrA binding-motifs (Gorla et al., 2018; Peterson et al., 2021)(**Supplemental Figure 5E**). We next confirmed that *mtrA* and *mtrB* knockdown led to a downregulation of putative MtrA regulon genes by RT-qPCR (**Supplemental Figure 5F**), validating the RNA-seq results. Surprisingly, in contrast to the results observed with *mtrAB* knockdown, silencing *lpqB* led to an upregulation of MtrA regulon genes (**Supplemental Figure 5G)**. In contrast to proposed role of LpqB as a positive regulator of this pathway (Nguyen et al., 2010), these data instead suggest that LpqB may be a negative regulator of MtrA signaling. To distinguish whether MtrA activation in the absence of LpqB requires MtrB, or whether loss of LpqB activates MtrA independent of MtrB, we next tested MtrA regulon expression upon simultaneous silencing of *mtrB* and *lpqB*. Consistent with the former model, MtrA activation required MtrB in the absence of LpqB (**Figure 2G**). These results, as well as the overlapping but distinct chemical-genetic interactions observed for *mtrAB* and *lpqB* (**Figure 2A**), are consistent with a model whereby the extracytoplasmic lipoprotein LpqB functions as a negative regulator of MtrB to restrain MtrA activation.

While the functions of most of the candidate MtrA regulon genes are unknown, a number of these genes encode peptidoglycan remodeling enzymes, including the endopeptidases *ripA*, *ripB*, *ripD*, the amidase *ami1*, and the transglycosylase *rpfC* (**Supplemental Data 3**) (Gorla et al., 2018; Peterson et al., 2021; Sharma et al., 2015). These gene products are important for proper peptidoglycan remodeling during growth and division. Intriguingly, a number of the MtrA regulon genes were also identified as sensitizing hits in the chemical-genetic screen, phenocopying the chemical-genetic effects of *mtrA* silencing (**Figure 2A**).

Transcript levels of neither *mtrAB* nor its regulon were altered in response to antibacterial challenge (**Supplemental Figure 5H**), indicating that unlike some two-component signaling systems in *Staphylococci* (Rajagopal et al., 2016; Yin et al., 2006), MtrAB is unlikely to be a stress-responsive signaling system.

Instead, these results highlight the central role of the MtrAB signal transduction pathway in coordinating proper peptidoglycan remodeling during bacterial growth and division (**Figure 2H**). These results further suggest that both inhibition (MtrAB inhibitors) or activation (LpqB inhibitors) of this pathway has the potential to prevent Mtb growth and dramatically reduce intrinsic drug resistance, highlighting the potential utility of small molecule inhibitors of this pathway.

### A diverse set of pathways contribute to intrinsic resistance and susceptibility to ribosome-targeting antibiotics

Unlike rifampicin, vancomycin, and bedaquiline, inhibition of mAGP biosynthesis did not sensitize Mtb to the three ribosome-targeting drugs streptomycin, clarithromycin, and linezolid (**Supplemental Figure 3C,D**; **Supplemental Data 1**). Thus, inhibition of mAGP biosynthesis is unlikely to be a relevant mechanism to potentiate the activity of these drugs. A prior publication reported that the cell-wall targeting drug ethambutol can synergize with clarithromycin (Bosne-David et al., 2000). In contrast to these results, neither our screen (**Supplemental Data 1**) nor checkerboard assays (**Supplemental Figure 6A**) validated a potentiating effect of ethambutol with clarithromycin, further validating the specificity of envelope-mediated intrinsic resistance for only a subset of antibiotics.

Streptomycin, the first successful antibiotic used to treat TB, is a natural product aminoglycoside which interacts with the 30S ribosomal subunit and induces mis-translation (**Figure 3A**) (Krause et al., 2016). Because streptomycin can be toxic it is currently reserved for the treatment of drug-resistant TB (Cohen et al., 2020). Clarithromycin is a semi-synthetic macrolide which targets the nascent polypeptide exit tunnel (NPET) and inhibits translation elongation in a sequence-specific manner (Kannan et al., 2014). Clarithromycin has minimal activity against Mtb both *in vitro* and *in vivo* (Luna-Herrera et al., 1995; Truffot-Pernot et al., 1995) and is used rarely as last resort, salvage therapy for multidrug-resistant TB (MDR-TB) (Seung et al., 2014). Linezolid is a synthetic oxazolidinone which targets the peptidyltransferase center (PTC) of the 50S ribosomal subunit, directly adjacent to the clarithromycin binding site, and also inhibits translation elongation in a sequence-specific manner (Marks et al., 2016). Linezolid is used as part of BPaL (Bedaquiline, Pretomanid, Linezolid), a new, potent combination therapy used to treat MDR-TB (Conradie et al., 2020).

**Figure 3:**
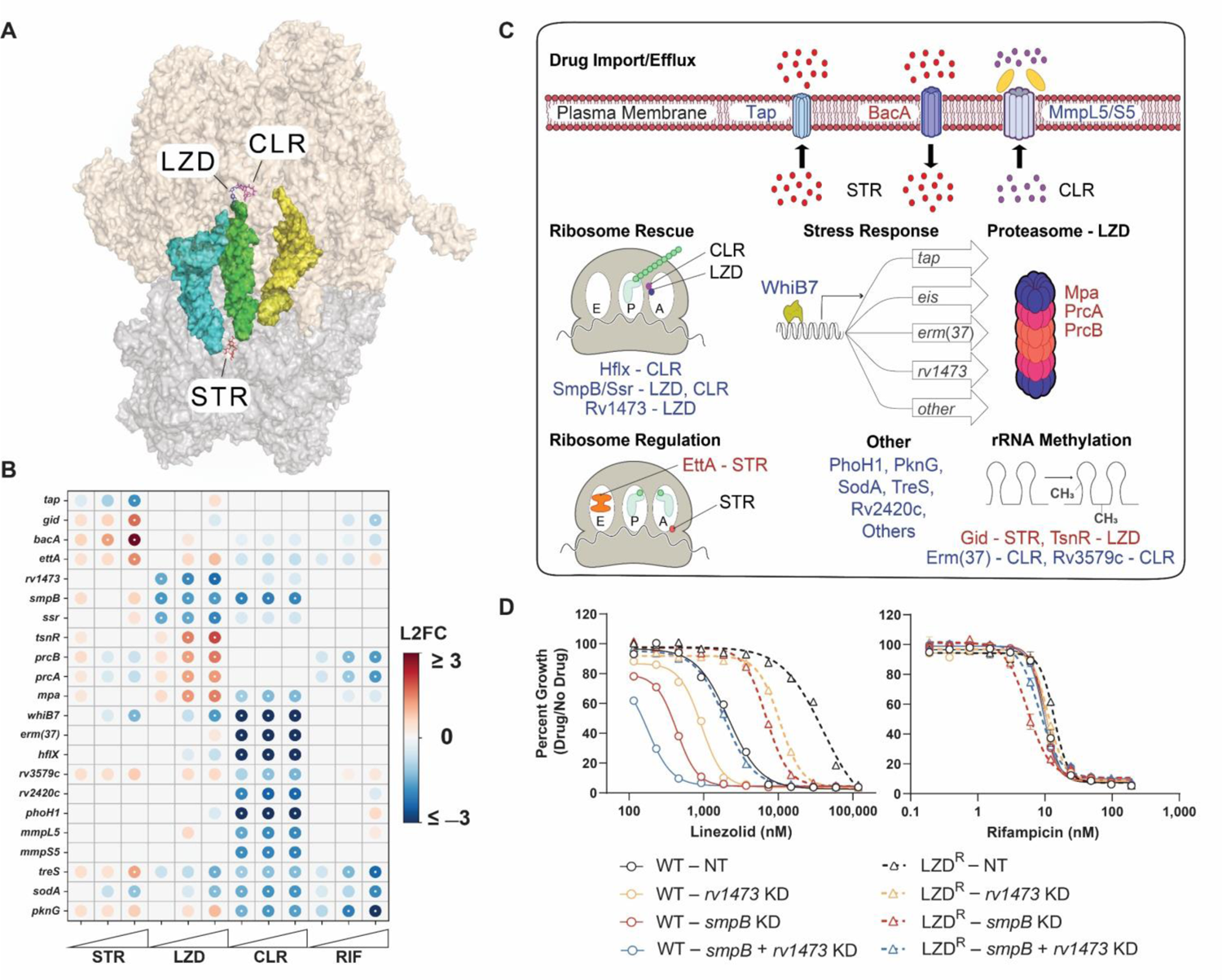
A diverse set of pathways contribute to intrinsic resistance and susceptibility to three ribosome targeting antibiotics in *M. tuberculosis*. (A) Structure of LZD (blue), CLR (magenta), and STR (red) bound to the ribosome of *Thermus thermophilus*. PDB codes: LZD (3DLL), CLR (1J5A), STR (1FJG), and ribosome with tRNAs (4V5C). (B) Feature-expression heatmap of select genes from the 5-day CRISPRi library pre-depletion screen. The color of each circle represents the gene-level L2FC; a white dot represents an FDR of < 0.01 and a |L2FC| > 1. (C) Chemical-genetic hit genes from panel (B) are involved in a diverse set of cellular pathways. Genes whose CRISPRi inhibition results in decreased or increased relative fitness in the presence of the three ribosome-targeting drugs are listed in blue or red font, respectively. (D) Single strain validation of LZD-associated hits. MIC values for LZD and RIF were measured for CRISPRi knockdown strains targeting *smpB* and *rv1473* in H37Rv or *rplC*-Cys154Arg linezolid-resistant H37Rv (LZD^R^). Error bars represent the standard error of the mean (SEM) for technical triplicates. Data are representative of at least two independent experiments.

Streptomycin, clarithromycin, and linezolid had correlated but distinct chemical-genetic signatures (**Figure 3B**, **Supplemental Figure 3B**). Clustering of the ribosome targeting drugs appeared to be driven in large part by the lack of an mAGP signature (**Supplemental Figure 3B,C; Supplemental Data 1)**, rather than any unique ribosome target signature, which may reflect the different mechanisms of action of the three ribosome targeting drugs. The sole sensitizing hit gene observed uniquely among the three ribosome targeting drugs was *whiB7*, a transcription factor that induces a stress response promoting intrinsic resistance to numerous ribosome-targeting antibiotics (Morris et al., 2005). Grouping hit genes based on predicted function connected the ribosome targeting antibiotics to both common and unique processes including: rRNA methylation, drug efflux/import, ribosome rescue, ribosome regulation, proteasome activity, and numerous poorly characterized genes (**Figure 3B,C**).

Consistent with prior publications, we found that rRNA methyltransferases can confer either intrinsic sensitivity or intrinsic resistance to ribosome-targeting drugs (Wilson, 2014). For example, silencing *erm(37)* resulted in strong depletion in clarithromycin treatment (**Figure 3B**), consistent with the role of Erm(37) in methylating the 23S rRNA to prevent macrolide binding (Madsen et al., 2005). Conversely, silencing the 16S rRNA methyltransferase *gid* resulted in strong enrichment in streptomycin treatment (**Figure 3B**), consistent with clinical observations whereby mutational inactivation of *gid* confers low-level acquired drug resistance to streptomycin (Wong et al., 2013). Interestingly, we found that knockdown of the predicted 23S rRNA methyltransferase *tsnR* confers resistance to linezolid (**Figure 3B**). This is analogous to work in *S. aureus*, in which loss of the evolutionarily distinct 23S methyltransferase *rlmN* confers linezolid resistance both *in vitro* and in the clinic (LaMarre et al., 2011; Pi et al., 2019). To determine if loss-of-function (LOF) mutations in *tsnR* could play a clinically relevant role in acquired linezolid resistance, we assembled a database of >45,000 whole genome sequences from Mtb clinical isolates (**Supplemental Data 4**). Given that linezolid has only recently been introduced to treat TB and is presently reserved to treat patients failing MDR-TB therapy, we expected linezolid-resistant TB to be rare. Consistent with this, we identified the most common linezolid-resistance promoting mutation *rplC*-Cys154Arg (Wasserman et al., 2019) only 122 times in our genome database. While putative LOF mutations in *tsnR* where even more rare (**Supplemental Data 5**), two MDR Mtb strains harbored both a *tsnR* frameshift allele (Thr156fs) and an *rplC*-Cys154Arg allele. The co-occurrence of these two mutations in two MDR Mtb strains is highly unlikely to have occurred by chance (𝜒^2^ test with Yates’ correction: p<0.0001). These data highlight that LOF *tsnR* mutations may serve as steppingstones to high-level resistance as linezolid is used more widely in the clinic.

Given the threat posed by linezolid resistance to future TB drug regimens and issues of linezolid toxicity, we next sought to determine if our findings could be exploited to identify synergistic drug-target combinations to overcome resistance and increase the therapeutic index for linezolid, analogous to pre-clinical efforts to boost ethionamide potency and tolerability (Blondiaux et al., 2017; Conradie et al., 2020). The essential trans-translation genes, *smpB* and *ssr* (Dejesus et al., 2017), were identified as strong linezolid sensitizing hits (**Figure 3B,C**). Trans-translation is a ribosome rescue pathway, thought to primarily rescue ribosomes stalled while translating non-stop mRNA transcripts (Alumasa et al., 2017). Due to its essentiality and potential importance for stress-tolerance, the trans-translation pathway has garnered attention as a mycobacterial drug target (Alumasa et al., 2017). Additionally, we identified the poorly characterized gene *rv1473* as a strong and specific linezolid sensitizing hit (**Figure 3B,C**). *rv1473* was previously reported to be a macrolide efflux pump (Duan et al., 2019). However, the lack of predicted transmembrane domains suggest *rv1473* is unlikely to be a membrane-embedded ABC transporter. Rather, homology suggests *rv1473* encodes an antibiotic resistance (ARE) ABC-F protein that functions to rescue linezolid-stalled ribosomes (**Supplemental Figure 6B**) (Antonelli et al., 2018). ARE ABC-F proteins bind to the ribosome to promote dissociation of ribosome-targeting antibiotics, thereby rescuing ribosomes from translation inhibition (Sharkey et al., 2016).

Our data suggest that inhibition of *rv1473* and trans-translation could make linezolid more potent, thereby lowering the dose of linezolid needed to inhibit bacterial growth. Confirming the screen predictions, individual CRISPRi knockdown of *rv1473* and *smpB* lowered the IC_50_ for linezolid by 2.3- and 5-fold respectively (**Figure 3D**; **Supplemental Figure 6C**) but did not alter sensitivity to other drugs. Inhibition of the Clp protease did not sensitize Mtb to linezolid (**Supplemental Figure 6D,E**), consistent with the critical role of trans-translation in rescuing linezolid-stalled ribosomes, not in Clp protease-mediated turnover of *ssrA*-tagged stalled translation products (Personne and Parish, 2014). Dual CRISPRi knockdown of both *rv1473* and *smpB* lowered the linezolid IC_50_ by 12.2-fold (**Figure 3D; Supplemental Figure 6C**), consistent with *rv1473* and trans-translation functioning in separate intrinsic resistance pathways. Interestingly, the increased linezolid sensitivity in the dual knockdown strain is similar to the magnitude of acquired resistance observed in linezolid-resistant clinical strains (Beckert et al., 2012). Thus, we hypothesized that dual inhibition of *rv1473* and *smpB* could functionally reverse linezolid resistance. Consistent with this hypothesis, dual knockdown of *rv1473* and *smpB* in a linezolid-resistant strain (*rplC*-Cys154Arg; **Supplemental Figure 6F**) restored linezolid sensitivity back to wild-type levels (**Figure 3D**; **Supplemental Figure 6G**), demonstrating that inhibition of intrinsic resistance factors can potentiate linezolid and functionally reverse acquired drug resistance.

### *bacA* mutations observed in Mtb clinical isolates confer acquired resistance to aminoglycosides and capreomycin

Acquired drug resistance is one of the greatest barriers to successful TB treatment. In recent decades, many acquired drug resistance mutations in Mtb have been mapped and characterized. However, our knowledge of the genetic basis of acquired drug resistance remains incomplete, particularly for mutations outside of the drug target or drug activator and which typically confer low to intermediate, but clinically relevant, levels of acquired drug resistance (Carter, 2021; Colangeli et al., 2018; Hicks et al., 2020; Walker et al., 2015). Given the ability of our chemical-genetic approach to identify hit genes associated with clinically relevant acquired drug resistance (**Figure 1B-D**, **Figure 3B**), we hypothesized that mining our chemical-genetic data may identify prevalent but previously unrecognized mechanisms of acquired drug resistance in Mtb.

We chose to focus our search for novel sources of acquired drug resistance to streptomycin. Streptomycin was introduced into the clinic in the late 1940s and remained an integral component of first-line TB therapy into the 1980s. It is now reserved to treat MDR-TB (Cohen et al., 2020). Unlike linezolid, which has only recently been used in the clinical management of TB, streptomycin has been used for almost eight decades. We hypothesized that this may have given rise to a diverse set of acquired resistance mutations which we could identify in our database of clinical Mtb genomes.

Aminoglycosides like streptomycin must traverse the Mtb envelope to access their ribosomal targets in the cytoplasm. The mechanism(s) by which aminoglycosides are taken up by mycobacteria are not well understood. Interestingly, and consistent with prior work (Domenech et al., 2009), the strongest hit gene leading to streptomycin resistance in our screen was *rv1819c* (*bacA*; **Figure 3B,C**). Recently, structural and biochemical work demonstrated that *bacA* is an ABC importer of diverse hydrophilic solutes (Rempel et al., 2020). Thus, we hypothesized that *bacA* may serve as an importer of streptomycin into the Mtb cytosol and that LOF mutations in *bacA* may be an unrecognized source of streptomycin resistance in clinical Mtb strains.

Searching our clinical strain genome database, we observed numerous *bacA* non-synonymous single nucleotide polymorphisms (SNPs) and small insertion-deletions (indels) and chose six for experimental validation (see *Materials and Methods* for more detail on SNP selection criteria; **Figure 4A**; **Supplemental Data 6**). Each mutation was introduced into a *bacA* expressing plasmid and transformed into a *bacA* deletion Mtb strain. Consistent with *bacA* LOF, four of the six alleles displayed an elevated streptomycin MIC (**Figure 4B**). Interestingly, these strains also showed elevated MICs to the other aminoyglycosides amikacin and kanamycin and the tuberactinomycin capreomycin but not to other drugs (**Figure 4B**; **Supplemental Figure 7A**). Moreover, overexpression of Mtb *bacA* in *M. smegmatis* sensitized *M. smegmatis* to streptomycin but not rifampicin or linezolid **(Supplemental Figure 7B)**. While further studies are necessary to definitively demonstrate that *bacA* is an importer of aminoglycosides and tuberactinomycins, our data, in combination with prior studies (Domenech et al., 2009; Rempel et al., 2020), strongly suggest that BacA imports these hydrophilic drugs **(Supplemental Figure 7C)** into the Mtb cytosol. Importantly, since a *bacA* deletion strain is not entirely resistant to aminoglycosides and tuberactinomycins, other relevant import mechanisms must exit in Mtb. These results demonstrate that our chemical genetic screens paired with clinical strain genomics can identify novel acquired drug resistance mutations in Mtb.

**Figure 4:**
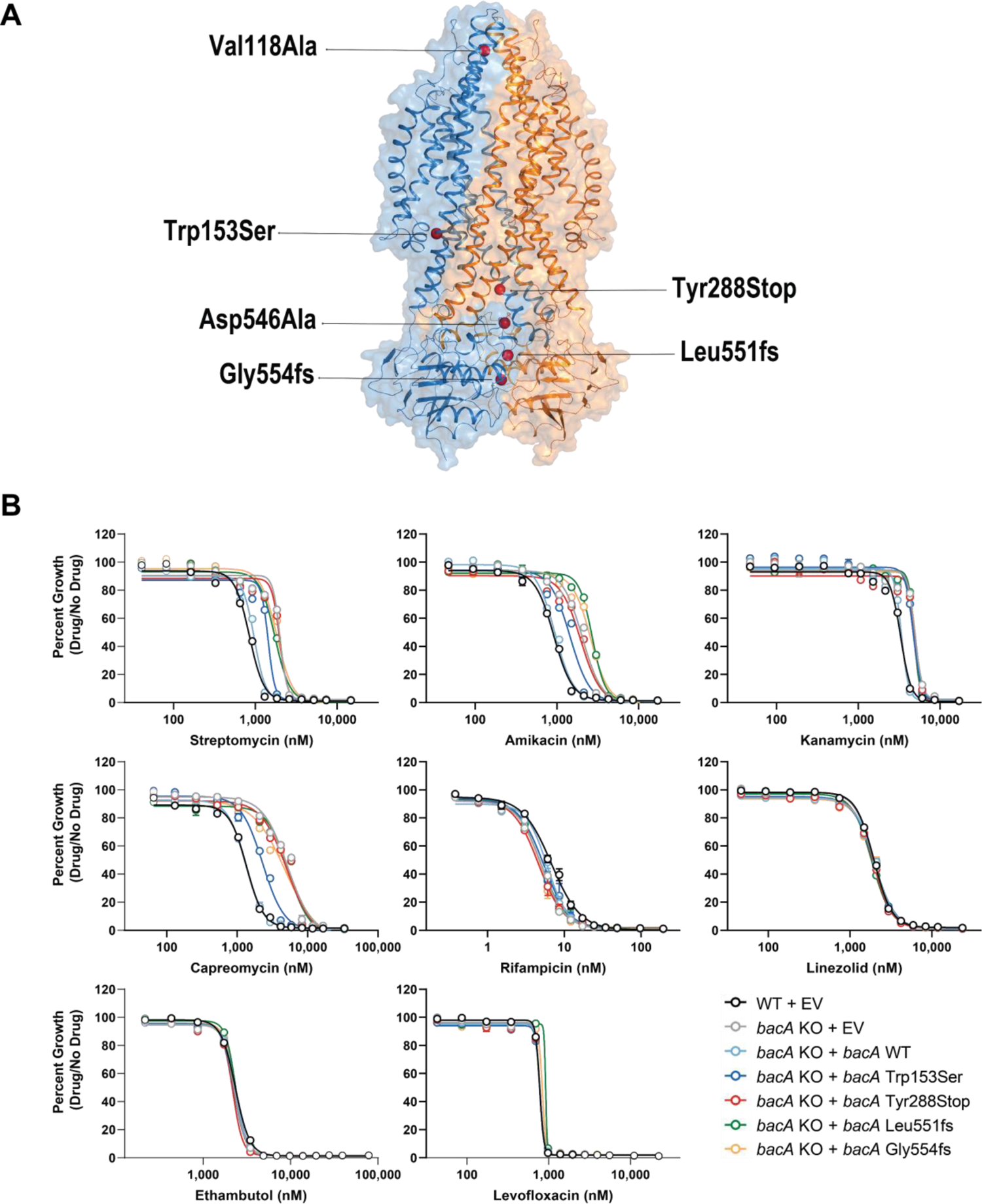
Loss-of-function mutations in *bacA* confer clinically relevant resistance to aminoglycosides and capreomycin. (A) Structure of *bacA* (PDB: 6TQF) (Rempel et al., 2020). Red spheres mark sites of experimentally tested clinical strain mutations in (C). (B) Drug resistance phenotypes for strains harboring *bacA* mutations. MIC values for the indicated drugs were measured for the indicated strains. Data represent mean ± SEM for technical triplicates. Results are representative data from at least two independent experiments.

### Partial loss-of-function mutations in *ettA* confer clinically relevant, low-level, acquired multidrug resistance

Another one of the strongest streptomycin resistance hits leading in our screen was *rv2477c*, which also showed low-level resistance to other drugs (**Figure 3B**; **Supplemental Data 1**). *rv2477c* is an ortholog of the *E. coli* gene *ettA* (∼58% amino acid identity), a ribosome-associated ABC-F protein that regulates the translation elongation cycle (Boël et al., 2014; Chen et al., 2013) (**Figure 3C, Figure 5A**). Due to its sequence similarity, we will refer to *rv2477c* as *ettA*. Biochemical studies demonstrated that ATP-bound EttA from *E. coli* stimulates formation of the first peptide bond of the initiating ribosome and then, concomitant with ATP hydrolysis, dissociates from the ribosome to allow translation elongation (Boël et al., 2014; Chen et al., 2013). Unlike *ettA* in *E. coli*, *ettA* is essential for the *in vitro* growth of Mtb (Bosch et al., 2021).

**Figure 5:**
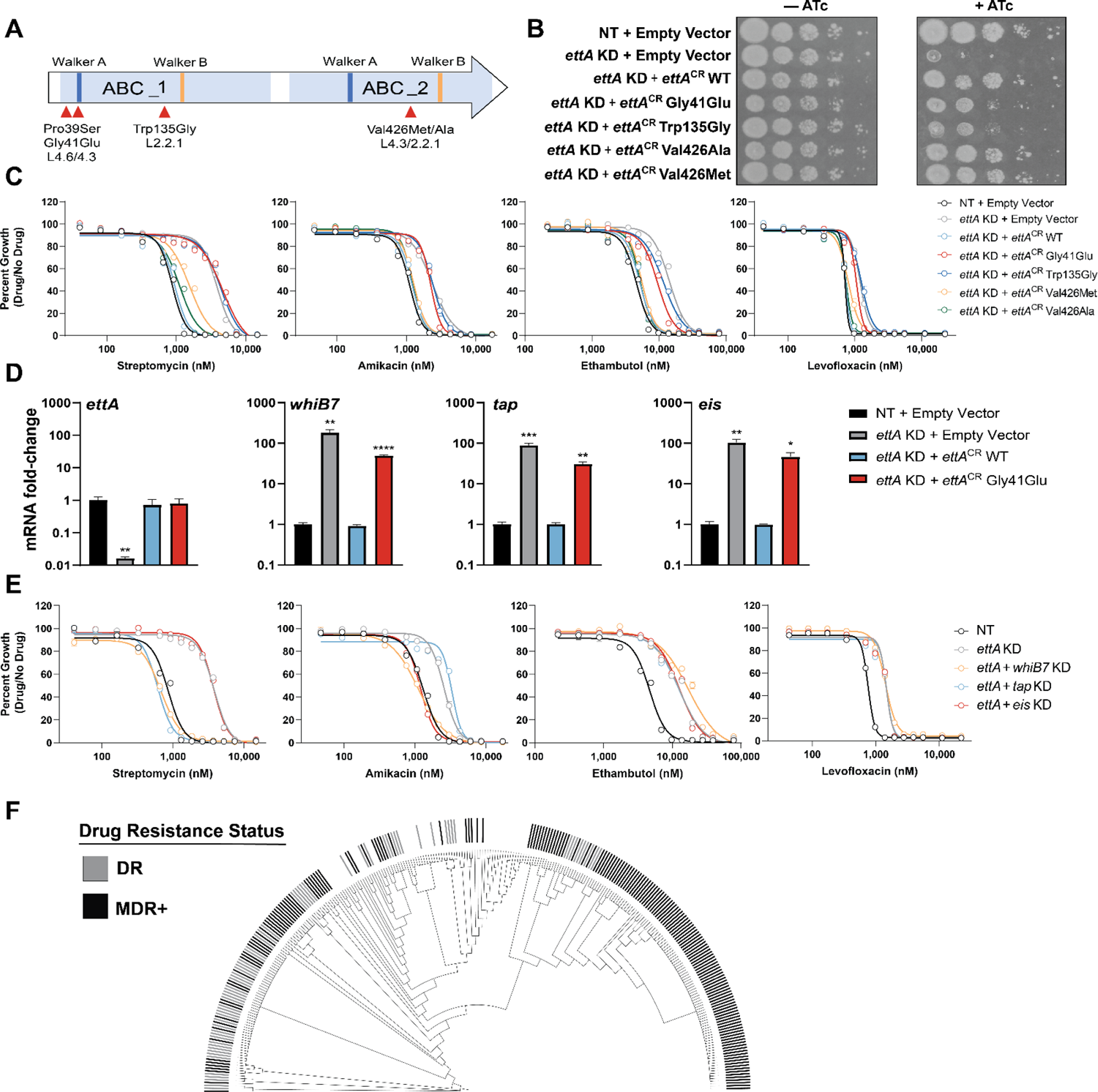
Partial loss-of-function mutations in *ettA* upregulate the *whiB7* stress response and confer low-level, acquired, multidrug resistance. (A) Domain organization of EttA. ABC domains are highlighted in light blue. Walker A and Walker B motifs are shown in dark blue and orange, respectively. SNPs observed in clinical Mtb isolates that were experimentally tested are highlighted with red arrows and the dominant lineage (L) in which that SNP is found is indicated. (B) Growth of *ettA* mutant and control strains. The *ettA* CRISPRi strain was complemented with an empty vector or CRISPRi-resistant alleles harboring the indicated SNPs. NT = non-targeting; WT = wild-type; KD = knockdown; CR = CRISPRi-resistant. (C) Drug resistance phenotypes for strains harboring *ettA* SNPs. MIC values for the indicated drugs were measured for the strains shown in **Figure 5B**. Data represent mean ± SEM for technical triplicates. Results are representative data from at least two independent experiments. (D) Quantification of indicated gene mRNA levels by qRT-PCR. Strains were grown in the presence of ATc for ∼5 generations prior to harvesting RNA. Error bars are SEM of three technical replicates. Statistical significance was calculated as p-value with Student’s t-test. *, p<0.05; **, p<0.01; ***, p<0.001, ****, p<0.0001. (E) MIC values for the indicated drugs were measured for *ettA* single and dual knockdown strains. Data represent mean ± SEM for technical triplicates. NT = non-targeting. (F) Phylogenetic tree of 291 Mtb clinical strains harboring the *ettA* Gly41Glu variant (**Supplemental Data 7**). Genotypically predicted drug-resistance status are shown. DR = resistance-conferring SNPs to RIF, INH, PZA, or EMB present; MDR+ = resistance-conferring SNPs to a minimum of RIF and INH.

Using our clinical Mtb strain genome database, we identified four non-synonymous SNPs as being located within motifs predicted to be important for EttA function (**Figure 5A**) and/or showed evidence for enrichment in genotypically predicted drug resistant Mtb strains (**Supplemental Data 7**). Lastly, we also included an *ettA* Trp135Gly mutation that was identified in serial Mtb isolates from a patient in Thailand. This Trp135Gly mutation was observed directly preceding the transition from MDR-TB to extensively drug-resistant (XDR) TB (Faksri et al., 2016). Candidate SNPs (**Figure 5A**) were incorporated into a CRISPRi-resistant *ettA* allele and transformed into an H37Rv strain that allowed selective CRISPRi silencing of the endogenous, wild-type *ettA* allele. All SNPs tested were capable of complementing knockdown of the endogenous *ettA* allele (**Figure 5B**). Both the Gly41Glu and Trp135Gly variants displayed a modest growth defect (**Figure 5B**; **Supplemental Figure 8A**), suggesting that these two SNPs are partial loss-of-function mutations.

Consistent with a role for these *ettA* SNPs in conferring acquired drug resistance, four out of the five variants showed an increased MIC for streptomycin (**Figure 5C**). The Gly41Glu and Trp135Gly strains showed a >5-fold shift in IC_50_, similar in magnitude to *gid* mutants (Wong et al., 2011). Additionally, the Gly41Glu and Trp135Gly mutants showed low-level resistance to a mechanistically diverse panel of antibiotics including amikacin, ethambutol, rifampicin, and levofloxacin, but not other tested drugs (**Figure 5C**; **Supplemental Figure 8C,D**).

To determine the mechanism by which *ettA* SNPs may confer low-level, acquired multidrug resistance, we analyzed the *M. smegmatis* proteome after silencing the *ettA* homolog, *ms4700*. Two of the most upregulated proteins upon *ms4700* knockdown were HflX (Ms2736) and Eis (Ms3513) (**Supplemental Figure 8E**) (Bosch et al., 2021), encoded by two genes known to be part of the *whiB7* regulon in Mtb (Morris et al., 2005). Thus, we hypothesized that partial loss of function *ettA* alleles may promote low-level, acquired multidrug resistance by stalling translation and constitutively upregulating the *whiB7* stress response– in essence, *ettA* mutations mimic the effects of translation stress caused by ribosome inhibitors to activate *whiB7* (Schrader et al., 2021). Consistent with this hypothesis, *whiB7* and known regulon genes (Morris et al., 2005) were constitutively upregulated in the *ettA* Gly41Glu Mtb mutant (**Figure 5D**).

Furthermore, simultaneous knockdown of *whiB7* was able to reverse aminoglycoside resistance conferred by knockdown of *ettA* (**Figure 5E**). This effect was drug-specific, with knockdown of the efflux pump *tap* specifically reversing streptomycin resistance and knockdown of the acetyltransferase *eis* specifically reversing amikacin resistance (Liu et al., 2019; Zaunbrecher et al., 2009). Interestingly, knockdown of *whiB7* did not reverse ethambutol or levofloxacin resistance, suggesting that the mechanism by which *ettA* mutations confer resistance to those drugs is independent of *whiB7*.

Further epidemiological analysis focused on *ettA* Gly41Glu,the most common *ettA* SNP in our database (n=291, ∼0.7% of all Mtb strains). Phylogenetic analysis shows that this cluster of related strains is heavily enriched for additional acquired drug resistance mutations (**Figure 5F, Supplemental Data 7**). Molecular epidemiology shows that *ettA* Gly41Glu strains are found in Spain, Italy, the United States (Couvin et al., 2019), but are concentrated in Peru (Sheen et al., 2013) and indigenous communities of Colombia (Marín et al., 2021), where they are driving a MDR-TB outbreak (**Figure 5F**).

### A loss-of-function mutation in *whiB7* renders an endemic Indo-Oceanic clade of *M. tuberculosis* hypersusceptible to macrolides

Since partial loss of *ettA* function appears to confer acquired drug resistance by constitutive activation of *whiB7*, we next mined our clinical strain genome database to identify putative gain-of-function *whiB7* mutations that may be associated with acquired drug resistance (Reeves et al., 2013). We identified numerous putative gain-of-function mutations in the *whiB7* promoter, 5’UTR, and upstream ORF (uORF), most of which have not been previously recognized as potential determinants of acquired drug resistance (**Figure 6A**; **Supplemental Data 7**) (Chakravorty et al., 2015; Kaur et al., 2016; Reeves et al., 2013).

**Figure 6:**
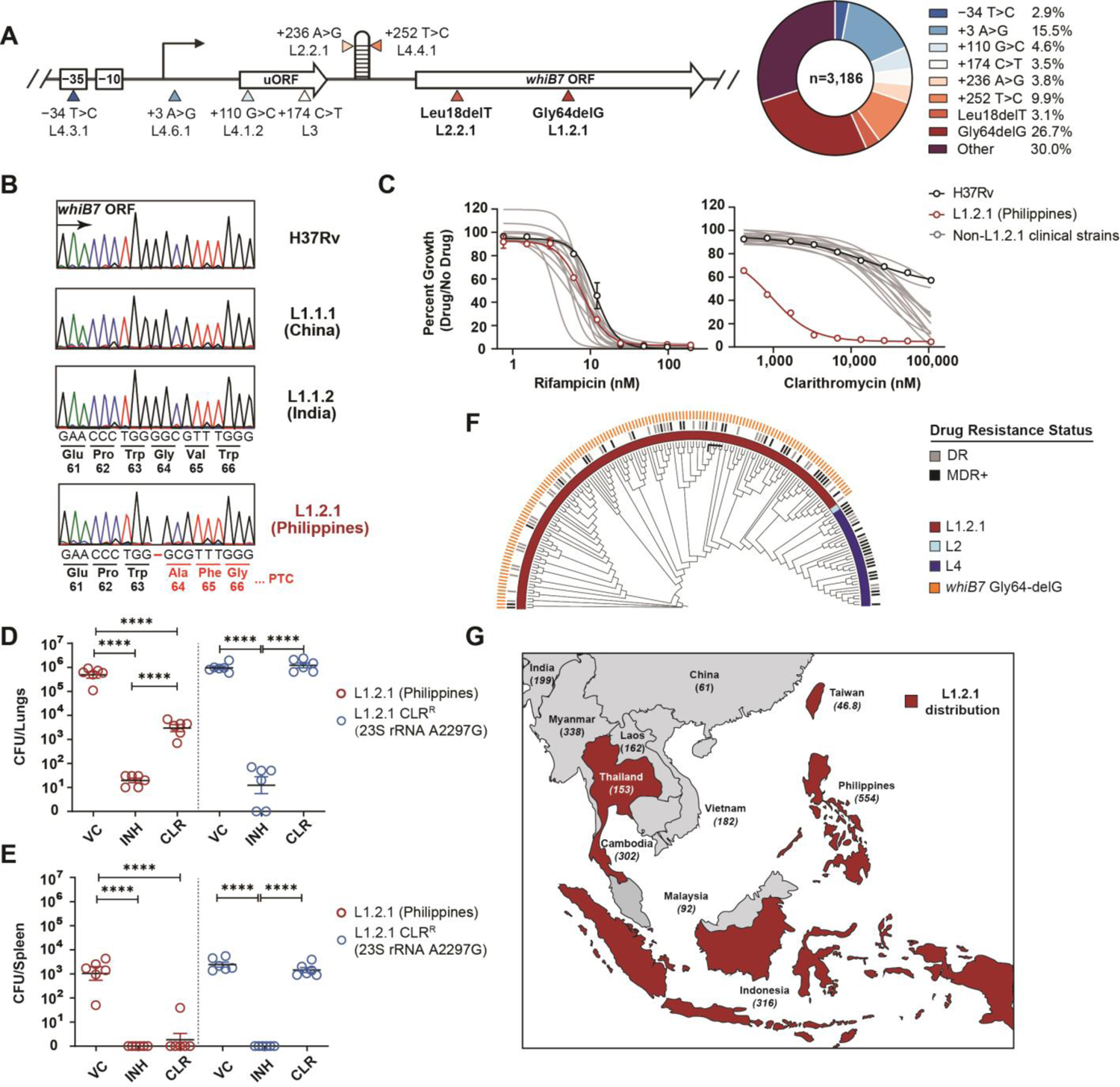
A loss-of-function mutation in *whiB7* renders an endemic Indo-Oceanic lineage of *M. tuberculosis* hypersusceptible to macrolides. (A) Diagram of Mtb *whiB7*. The eight most common *whiB7* variants observed in our Mtb clinical strain genome database are highlighted with arrows and the dominant lineage (L) in which those SNPs are found is indicated. The pie chart depicts the observed frequencies of each indicated variant in our database. (B) Sanger sequencing traces of *whiB7* from the indicated Mtb clinical strains and their country of origin. PTC = premature termination codon. (C) MIC values for RIF and CLR were measured for a reference set of Mtb clinical strains. Error bars represent the standard error of the mean (SEM) for technical triplicates. Results are representative data from at least two independent experiments. (D,E) Mean lung (D) and spleen (E) Mtb CFU (± SEM) in BALB/c mice after isoniazid (INH; 25 mg/kg), or clarithromycin (CLR; 200 mg/kg) treatment. Mice were infected with approximately 100-200 CFU of aerosolized Mtb. After 10 days to allow the acute infection to establish, chemotherapy was initiated. Following 24 days of drug therapy, Mtb bacterial load of lungs and spleen were determined. Statistical significance was assessed by one-way ANOVA followed by Tukey’s post-hoc test. ****, p<0.0001. VC = vehicle control. CLR^R^ = clarithromycin-resistant. (F) Phylogenetic tree of 178 Mtb clinical strains isolated during the 2012 nationwide drug resistance survey in the Philippines (Phelan et al., 2019) (**Supplemental Data 8**). The presence of the *whiB7* Gly64delG mutation and genotypically predicted drug resistance status are shown. DR = resistance-conferring SNPs to RIF, INH, PZA, or EMB present; MDR+ = resistance-conferring SNPs to a minimum of RIF and INH. (G) Map showing the distribution of the L1.2.1 sub-lineage in Southeast Asia (WHO, 2021). Tuberculosis incidence rates are listed in parentheses beneath each country name.

Unexpectedly, however, the most common *whiB7* variant in our database was a putative loss-of-function allele. This allele, Gly64delG, harbors a single nucleotide deletion at codon Gly64 and represents nearly one-third (n=851/3,186) of all *whiB7* variants in our database (**Figure 6A**) (Merker et al., 2020; Vargas et al., 2021; Warit, 2015). The Gly64delG frameshift results in a premature stop codon and truncation of the critical DNA binding AT-hook element (**Supplemental Figure 9A**) (Burian et al., 2013), thus presumably rendering WhiB7 inactive in these strains. *whiB7*-mediated intrinsic drug resistance typically renders macrolides ineffective to treat TB. We next sought to explore the possibility that the common Gly64delG mutation may render this subset of Mtb strains hypersusceptible to and treatable with macrolides.

Lineage calling identified the Gly64delG SNP as uniquely present in all lineage 1.2.1 (L1.2.1) Mtb isolates, a major sublineage of the L1 Indo-Oceanic clade (**Figure 6A**; **Supplemental Figure 9B**; **Supplemental Data 8)** (Netikul et al., 2021). Using a reference set of Mtb clinical strains (Borrell et al., 2019), we first validated the presence of the *whiB7* Gly64delG frameshift mutation in L1.2.1 by Sanger sequencing (**Figure 6B**). All other clinical isolates in this set were wild-type for *whiB7*. Consistent with loss of *whiB7* function, the L1.2.1 isolate was hypersusceptible to clarithromycin as well as other macrolides, ketolides, and lincosamides, whereas all other clinical isolates were intrinsically resistant (**Figure 6C; Supplemental Figure 9C-E**). The *whiB7* Gly64delG allele failed to complement intrinsic clarithromycin resistance in an H37Rv Δ*whiB7* knockout strain, confirming that Gly64delG is a loss-of-function allele (**Supplemental Figure 9F**). To confirm L1.2.1 macrolide susceptibility *in vivo*, we infected mice with H37Rv or L1.2.1 Mtb by low-dose aerosol exposure. Both strains showed similar growth kinetics *in vivo* (**Supplemental Figure 10A,B**). We next tested drug efficacy in an acute infection model, with drug dosing designed to mimic human pharmacokinetics in the treatment of TB (rifampicin, isoniazid) and non-tuberculous mycobacteria (clarithromycin) (Rodvold, 1999). We isolated a spontaneous clarithromycin resistant L1.2.1 isolate (harboring a 23S A2297G mutation) as a control (**Supplemental Figure 10C-E**). Consistent with the *in vitro* data, L1.2.1 was sensitive to macrolide therapy whereas H37Rv was intrinsically resistant (**Figure 6D,E; Supplemental Figure 10F-I**) (Luna-Herrera et al., 1995; Truffot-Pernot et al., 1995). Therapeutic drug monitoring confirmed equivalent drug exposures in both the H37Rv and L1.2.1 infections (**Supplemental Figure 10J,K**). Interestingly, L1.2.1 was also more sensitive than H37Rv to rifampicin in the mouse model (**Supplemental Figure 10H,I**), which could reflect *whiB7*-dependent upregulation of the tap efflux pump in H37Rv but not L1.2.1 during induced tolerance *in vivo* (Adams et al., 2011)

To estimate the potential clinical impact of this finding, we next examined the geographic distribution of the L1.2.1 sublineage. This sublineage is found predominantly in Southeast Asia, including the high TB burden countries Indonesia, Thailand, and the Philippines (**Figure 6F,G**) (Palittapongarnpim et al., 2018). L1.2.1 is particularly prevalent in the Philippines, accounting for approximately 80% of all Mtb isolates in this country (Phelan et al., 2019). Of note, the Philippines has one of the highest TB incidence rates in the world, including a high burden of drug-resistant TB, and TB is a leading cause of death in this country (WHO, 2021). A recent analysis of the global burden of TB caused by L1.2.1 estimates that this sublineage causes approximately 600,000 cases of active TB per year (Netikul et al., 2021), of which ∼43,000 are estimated to be MDR-TB based on the frequencies of drug resistance in our clinical strain genome database. Thus, clarithromycin, an effective, orally available, safe, and generic antibiotic, could potentially be repurposed to treat a major sublineage of TB.

## DISCUSSION

A deeper understanding of the bacterial pathways that govern drug efficacy in Mtb is needed to develop more potent therapies, identify new mechanisms of acquired drug resistance, and reveal overlooked therapeutic opportunities. To address this challenge, we developed a CRISPRi platform to define the genetic determinants that alter bacterial fitness in the presence of different drugs, and then overlay these chemical-genetic results with comparative genomics of Mtb clinical isolates. Illustrating the power of this dataset to derive new, clinically relevant biological insight, we uncover diverse mechanisms of intrinsic drug resistance that can be targeted to potentiate therapy, describe new mechanisms of acquired drug resistance associated with the emergence of MDR-TB, and make the unexpected discovery of an “acquired drug sensitivity” that could enable the repurposing of clarithromycin to treat an Mtb sublineage.

Alternative functional-genomics methods such as TnSeq have been successfully applied to generate chemical-genetic interaction profiles for a number of drugs in Mtb (Sassetti et al., 2020; Xu et al., 2017). While powerful, TnSeq as currently implemented in Mtb is restricted to the analysis of *in vitro* non-essential genes and thus cannot assess some of the most compelling drug targets, essential genes. The recent development of barcoded degron libraries in Mtb, in which regulated proteolysis is used to tune target protein levels, now allows the chemical-genetic assessment of hundreds of essential genes (Johnson et al., 2019; Koh et al., 2021). This approach is likely to be expanded to include nearly all essential genes in the near future. The degron library has been used to identify numerous new inhibitor-target pairs in large-scale, target-based whole cell screens. The degron approach is extremely powerful but suffers from the fact that not all proteins tolerate the degron tag, the approach fails to assess chemical-genetic interactions for non-essential genes, and the laborious nature of mutant construction functionally restricts analysis to a single Mtb reference strain (H37Rv). By being able to robustly tune knockdown for both essential and non-essential genes, the CRISPRi approach taken here provides the most comprehensive chemical-genetic map available, successfully generating distinct profiles for two translation inhibitors that bind within angstroms of each other. Moreover, the portability of CRISPRi libraries will allow chemical-genetic screening across diverse Mtb clinical isolates (Bosch et al., 2021). While more comprehensive and portable, our current CRISPRi approach is low throughput compared to degron libraries, and thus the development of optimized, compact CRISPRi libraries to increase screen throughput remains a priority. CRISPRi has well-known limitations (Bosch et al., 2021), including the polar effect of CRISPRi knockdown. As with any genetic approach, genetic inhibition of a target is not the same as inhibition of a target with a small molecule (Knight and Shokat, 2007), highlighting the importance of validating chemical-genetic interactions with small molecule inhibitors (Cokol et al., 2017). Lastly, any pooled screening approach may miss effects where the phenotype can be complemented *in trans* (e.g. cross-feeding), although recent TnSeq results suggest that only 1-3% of all *S. pneumoniae* Tn mutants show differential fitness phenotypes when grown as pools vs in isolation, suggesting this type of phenotypic masking is rare (Thibault et al., 2019).

One proposed route to improving TB chemotherapy is to develop more potent drug combinations by leveraging drug synergies (Cokol et al., 2017). By identifying hundreds of genes that contribute to intrinsic drug resistance in Mtb, the results presented here can be used to inform drug development efforts to identify synergistic drug combinations that disarm intrinsic drug resistance. Our results confirm that one of the richest sources of potentially synergistic targets is the mycobacterial envelope, consistent with the long-appreciated understanding of the envelope as a barrier to antibiotic efficacy (Batt et al., 2020; Jarlier and Nikaido, 1994; Xu et al., 2017). mAGP disruption may increase envelope permeability and antibacterial uptake (McNeil et al., 2019; Piddock et al., 2000), as confirmed for *kasA* and *mtrAB*; alternatively, the chemical-genetic results may indicate a more mechanism-specific interaction, whereby knockdown of mAGP biosynthetic or regulatory genes is synthetic lethal with subinhibitory concentrations of an antibacterial compound (Xu et al., 2017). It is tempting to speculate that potentiation of rifampicin could, at least in part, explain the clinical success of ethambutol. Long-thought to be included in the standard regimen primarily to minimize the emergence of drug resistance to isoniazid, pyrazinamide, and rifampicin (Zimmerman et al., 2017), ethambutol may have “won” in early clinical trials due to its ability to effectively penetrate lung lesions (Zimmerman et al., 2017) and potentiate rifampicin (Cokol et al., 2017; Piddock et al., 2000). Chemical-genetic profiling of an expanded set of antibacterials to identify those compounds potentiated by mAGP disruption, or other types of molecules for which uptake can be monitored, may allow derivation of the permeability “rules” of the Mtb cell envelope, which could then be used to guide drug development to increase compound permeability (Davis et al., 2014).

Beyond the Mtb cell envelope, our results uncover both shared and unique intrinsic resistance and sensitivity mechanisms, as highlighted by the chemical-genetic profiles for the three ribosome targeting antibiotics. Future biochemical studies will identify the molecular mechanisms by which these factors operate, knowledge which could then be used to guide medicinal chemistry efforts to improve these drugs. For example, aminoglycoside scaffolds could be designed to improve BacA-mediated uptake (Domenech et al., 2009; Rempel et al., 2020). Similarly, our results suggest that Rv1473 serves as an antibiotic resistance ABC-F protein, capable of displacing oxazolidinones and phenicols from the Mtb ribosome (**Supplemental Figure 6**). Next generation oxazolidinone analogs could be designed that are recalcitrant to the potential drug-displacing activity of Rv1473, analogous to the third-generation tetracycline analogues which are resistant to the drug-displacing activity of the ABC-F protein TetM (Jenner et al., 2013). In light of our findings, we suggest designating *rv1473* as *oprA* (**<u>o</u>**xazolidinone **<u>p</u>**henicol **<u>r</u>**esistance **<u>A</u>**). Finally, these results show that trans-translation may be a target for synergistic drug combinations (Brunel et al., 2018), which could be important in increasing the potency and decreasing toxicity of oxazolidinones.

In addition to guiding rational development of synergistic drug combinations, our results illustrate the power of combining chemical-genetics with comparative genomics to discover new mechanisms of acquired drug resistance. In recent decades, many drug resistance mutations in Mtb have been mapped and characterized. However, our knowledge of the genetic basis of acquired drug resistance remains incomplete, particularly for mutations outside of the drug target or drug activator and which typically confer low to intermediate levels of drug resistance (Carter, 2021; Hicks et al., 2018; Walker et al., 2015). Low-level drug resistance has been associated with TB treatment failure (Colangeli et al., 2018), and could serve as a stepping stone to allow additional, high-level drug resistance mutations to evolve (Dick and Dartois, 2018). We make a number of findings that may be important for diagnosing and treating drug resistant TB. First, we show inhibition of *tsnR* increases fitness in the presence of linezolid, and thus mutations in this gene could be monitored as linezolid use is expanded in the clinic. Second, we find that LOF mutations in the ABC importer *bacA* confer resistance to four important second-line TB drugs: streptomycin, kanamycin, amikacin, and capreomycin. These mutations may be a source of unexplained resistance amongst clinical Mtb strains. Third, we show that partial loss-of-function mutations in the essential gene *ettA* result in constitutive activation of the *whiB7* stress response and low-level acquired multidrug resistance. This phenotype is consistent with the TB patient described in (Faksri et al., 2016), whereby serial Mtb isolates acquired a Trp135Gly mutation in *ettA* directly preceding the transition from MDR-TB to extensively drug-resistant XDR-TB. 3.1% (n=1,393/45,473) of all Mtb strains in our genome database harbor a missense SNP in *ettA*, suggesting that *ettA*-mediated acquired drug resistance could be highly prevalent. Our focus on the *ettA*-Gly41Glu mutation shows that it is highly prevalent and that it likely facilitated the evolution of an MDR-TB outbreak concentrated in Peru, a country with one of the highest MDR-TB burdens in South America (WHO, 2021). Our results demonstrate that while streptomycin and amikacin may be less effective against *ettA* variants, capreomycin may remain effective and should be considered for treatment. In addition to *bacA* and *ettA*, our analytical pipeline revealed numerous additional genes as candidates for previously unrecognized mechanisms of acquired drug resistance in Mtb (**Supplemental Data 9**), although further work is necessary to validate these predictions. An increased understanding of acquired drug resistance will guide development of more effective molecular diagnostics and personalized TB therapy to reduce treatment failure and the subsequent evolution of additional resistance alleles.

In the search for gain-of-function *whiB7* mutations that confer acquired drug resistance, we made the unexpected discovery of common loss-of-function *whiB7* alleles (Merker et al., 2020; Vargas et al., 2021), which we refer to as an “acquired drug sensitivity.” In addition to *whiB7*, we identified several predicted loss-of-function alleles in other genes that could confer acquired drug sensitivity in other Mtb clinical strains (**Supplementary Table 2**), and validate LOF alleles in a L1 and L7 isolate that confer hypersusceptibility to bedaquiline and the anti-leprosy drug clofazimine (**Supplemental Figure 11**) (Carter, 2021). Phylogenetic dating suggests that the *whiB7* mutation arose approximately 900 years ago, well before the introduction of TB chemotherapy (O’Neill et al., 2019). Since macrolides and lincosamides have not historically been used to treat TB, there has likely been little selective pressure against *whiB7* loss-of-function mutants. Whether the Gly64delG mutation provides or provided a selective benefit to L1.2.1 in Southeast Asia, enriched as a passenger mutation due to strong linkage disequilibrium in Mtb, enriched as a result of epistatic interactions with the L1.2.1 genotype that negate the selective benefit of the *whiB7* stress response, or was simply the product of genetic drift remains unclear. We find that the entire L1.2.1 Mtb sublineage (Merker et al., 2020) is a *whiB7* loss-of-function mutant which renders this strain susceptible to macrolides, both *in vitro* and *in vivo*. L1.2.1 could be identified by molecular diagnostics such as Genexpert (Walker et al., 2015). These results pave the way for further preclinical efficacy studies to support that clarithromycin be repurposed to treat this major Mtb sublineage (∼600,000 active TB cases per year, ∼43,000 MDR TB cases, 80% of all TB in the Philippines) in Southeast Asia.

In summary, we combine genome-scale CRISPRi chemical-genetics and comparative genomics of Mtb clinical strains to define bacterial mechanisms that limit drug efficacy. This chemical-genetic map provides a rich resource to guide development of more potent drugs and drug combinations, identify previously unrecognized mechanisms of acquired drug resistance, and highlights overlooked therapeutic opportunities. Chemical-genetic profiling of antibacterials not traditionally used to treat TB may identify additional acquired drug sensitivities that could be leveraged to repurpose such drugs to treat TB. Profiling of lead compounds early in drug discovery, in addition to providing (or refuting) evidence for on-target activity (**Figure 1B-D**), may allow the identification of relevant bacterial intrinsic resistance mechanisms, knowledge which could then be used to modify the leads to evade intrinsic resistance (Lee et al., 2014). Future iterations of this approach should address additional bacterial mechanisms that contribute to treatment failure, including drug tolerance and persistence (Hicks et al., 2018). Moreover, it is well appreciated that chemical-genetic interactions can be strongly influenced by genetic background and growth environment (Bosch et al., 2021; Koh et al., 2021). This work sets the stage for expanded chemical-genetic studies in different Mtb clinical strains and different growth environments, including *in vivo* infection models.

## ACKNOWLEDGEMENTS

We thank members of the Rock laboratory, Sabine Ehrt, Justin Pritchard, Alexandre Gouzy and Shipra Grover for comments on the manuscript and/or helpful discussions. We thank Carolina Trujillo, Josh Wallach, Sophie Lavalette-Levi, and Sabine Ehrt for technical assistance, Jenny Xiang and Dong Xu of the Weill Cornell Genomics Core for sequencing, the animal technical and analytical teams of the Center for Discovery and Innovation for animal work, and Sarah Schrader for sharing the *M. smegmatis whiB7* knockout strain. We thank Ruby Froom and Viviana LaSalle for scientific illustration. We thank Cliff Barry for kindly sharing the *bacA* deletion strain used in this work. This work was supported by a Harvey L. Karp Postdoctoral Fellowship (S.L.), the Potts Memorial Foundation (M.D. and S.L.), the Bill and Melinda Gates Foundation (INV-010616 and INV-004761 to D.S.), the Department of Defense (PR192421 to J.R.), an NIH Shared Instrumentation Grant (S10-OD023524 to V.D.), and an NIH/NIAID New Innovator Award (1DP2AI144850-01, J.R.).

## AUTHOR CONTRIBUTIONS

Conceptualization, S.L., N.C.P., N.R., D.S. and J.M.R.; Methodology, S.L., N.C.P., J.S.C., Z.A.A., M.A.D., and J.M.R.; Investigation, S.L., N.C.P., N.R., M.D.Z., B.B., C. E., D.S., K.P., M.G. and K.R.; Validation: S.L. and N.C.P.; Software & Formal Analysis: J.S.C., M.A.D., Z.A.A. and K.E.; Data Curation: J.S.C., M.A.D., Z.A.A., K.E. and J.M.R.; Writing – Original Draft, S.L., N.C.P. and J.M.R.; Writing – Review & Editing, S.L., N.C.P., J.S.C., M.A.D., K.E., B.B., M.G., V.D., D.S. and J.M.R.; Funding Acquisition, V.D., D.S. and J.M.R.; Resources, M.D.Z., M.G., V.D.; Supervision, J.M.R.

## Supplemental Figures

**Supplemental Figure 1:**
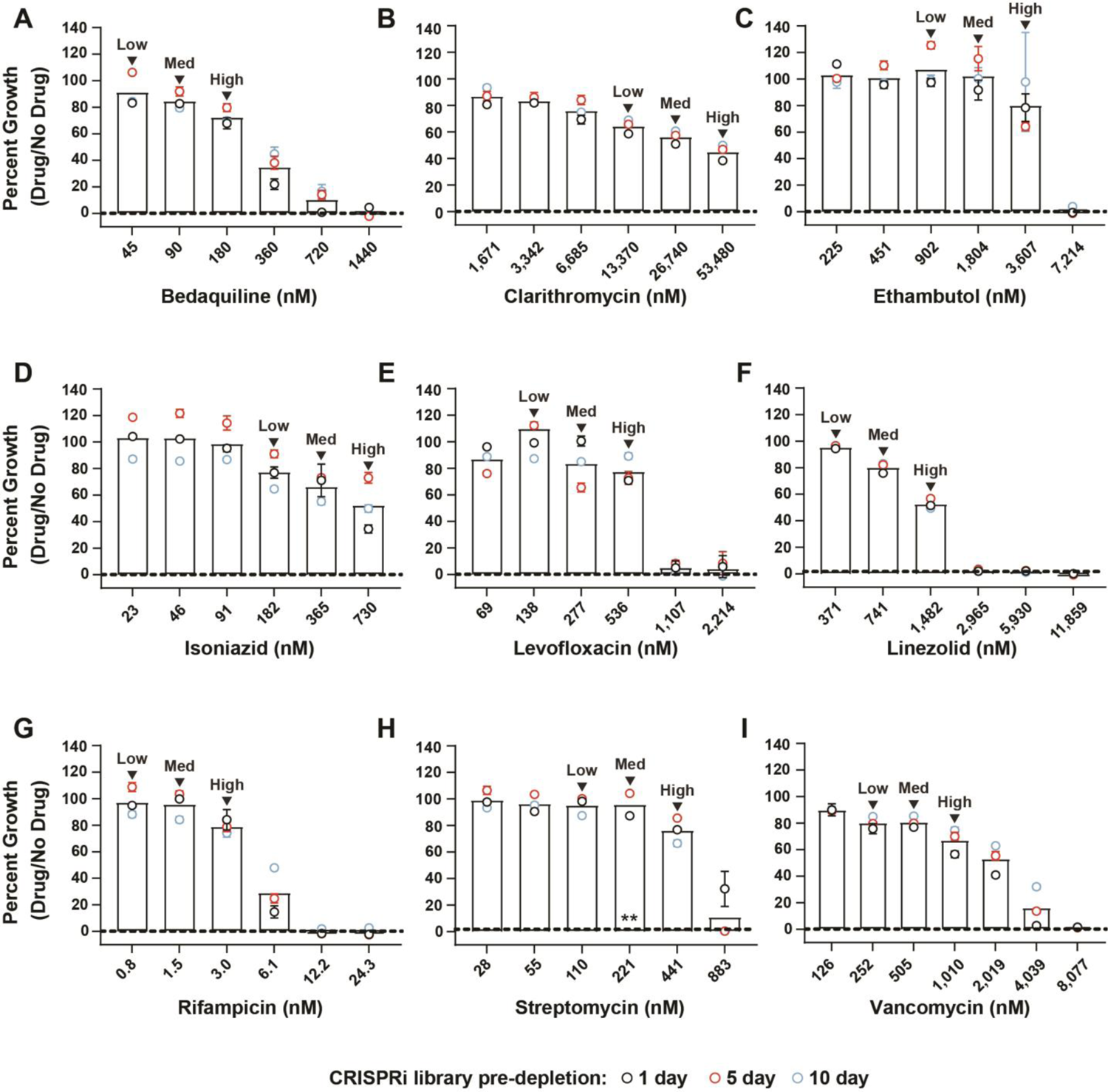
Growth of the Mtb CRISPRi library during drug selection. (A-I) Normalized growth of the Mtb CRISPRi library in the drug screens. Error bars represent the SEM for biological triplicates. Samples harvested for sgRNA deep sequencing are marked as “High”, “Med”, and “Low”, denoting the three descending doses of partially inhibitory drug concentrations analyzed in these screens. **: 10-day sample was lost for the 221 nM (“Med”) streptomycin screen.

**Supplemental Figure 2:**
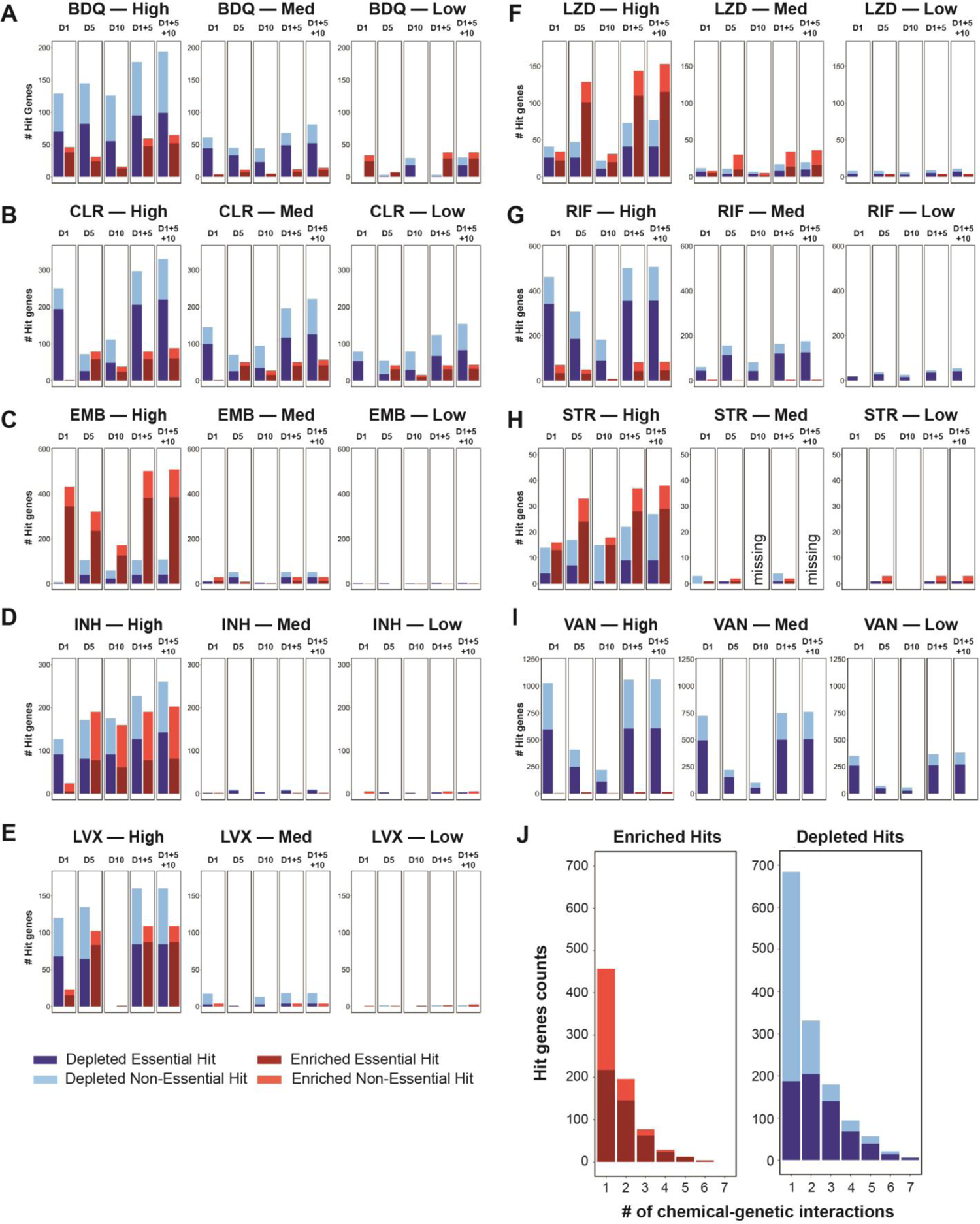
Summary of hits from chemical-genetic screens. (A-I) Bar graphs showing number of hit genes identified across all conditions. Gene essentiality calls were defined by CRISPRi as in (Bosch et al., 2021). D1, D5, and D10 indicate the number of days the CRISPRi library was treated with ATc prior to drug exposure; D1+5 = hit genes defined as the union of 1 and 5-day CRISPRi library pre-depletion results; D1+5+10: hit genes defined as the union of 1, 5 and 10 day CRISPRi library pre-depletion results. Note that the 10-day sample was lost for the “Med” streptomycin screen and thus the D10 containing results for “STR–Med” are labelled “missing”. (J) Histogram depicting the number of unique chemical-genetic interactions for enriching and depleting hits. Hit genes were defined as the union of 1 and 5-day CRISPRi library pre-depletion results.

**Supplemental Figure 3:**
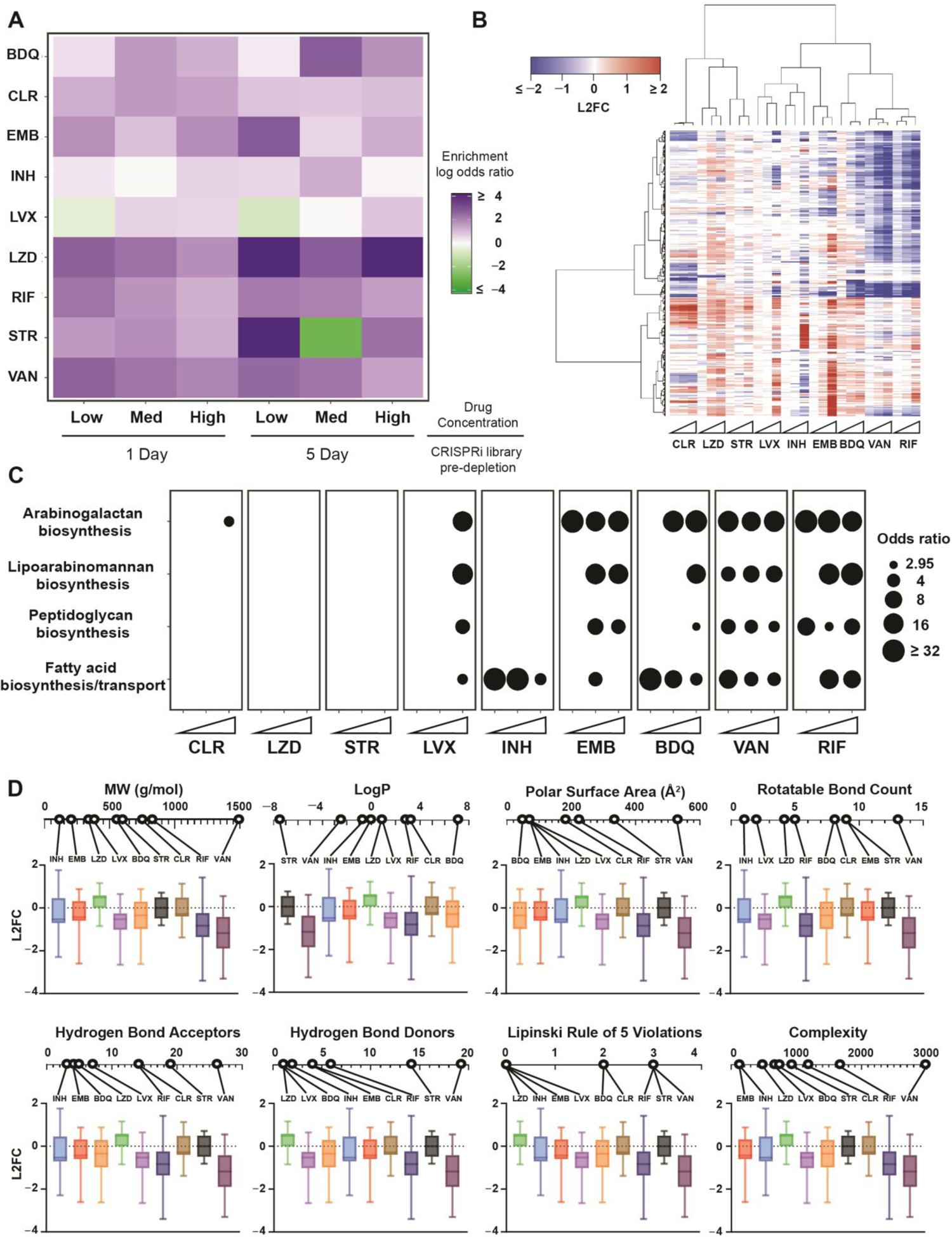
Clustering & enrichment analysis of chemical-genetic profiles. (A) Heatmap of odds-ratios showing enrichment of essential gene targeting sgRNAs as hits in the chemical-genetic screen. A Fisher exact test was used to evaluate enrichment of essential gene targeting sgRNAs relative to non-essential gene targeting sgRNAs amongst hit genes (FDR <0.01, |L2FC| > 1) in the chemical genetic screen. (B) Heatmap showing clustered chemical-genetic profiles from the 5-day CRISPRi library pre-depletion screen. Genes are clustered along the vertical axis; for simplicity, only genes that hit in at least two drugs are shown (n=676 genes). Ascending drug concentrations (“Low”, “Med”, “High” indicated by white triangles) are clustered along the horizontal axis. The median L2FC for each gene following drug selection (relative to vehicle control) is indicated on the color scale. If a gene was not a significant hit (FDR > 0.01), the L2FC value was plotted as 0 for the corresponding condition. The full dataset is available in **Supplemental Data 1**. (C) Bubble plot of the enriched (P < 0.05) KEGG categories for hit genes for the indicated drugs. KEGG annotations were manually updated to include the mycolic acid-arabinogalactan-peptidoglycan (mAGP) complex-associated genes described in (Jankute et al., 2015; Maitra et al., 2019). (D) Correlation of mAGP signature and physiochemical properties. For each drug, the distribution of L2FC values (“High” concentration, 5-day CRISPRi library pre-depletion) is shown for a select group of 78 genes involved in mAGP assembly and regulation as described in (Jankute et al., 2015; Maitra et al., 2019). In each plot the drugs are arranged based on their numerical value for each given physiochemical property.

**Supplemental Figure 4:**
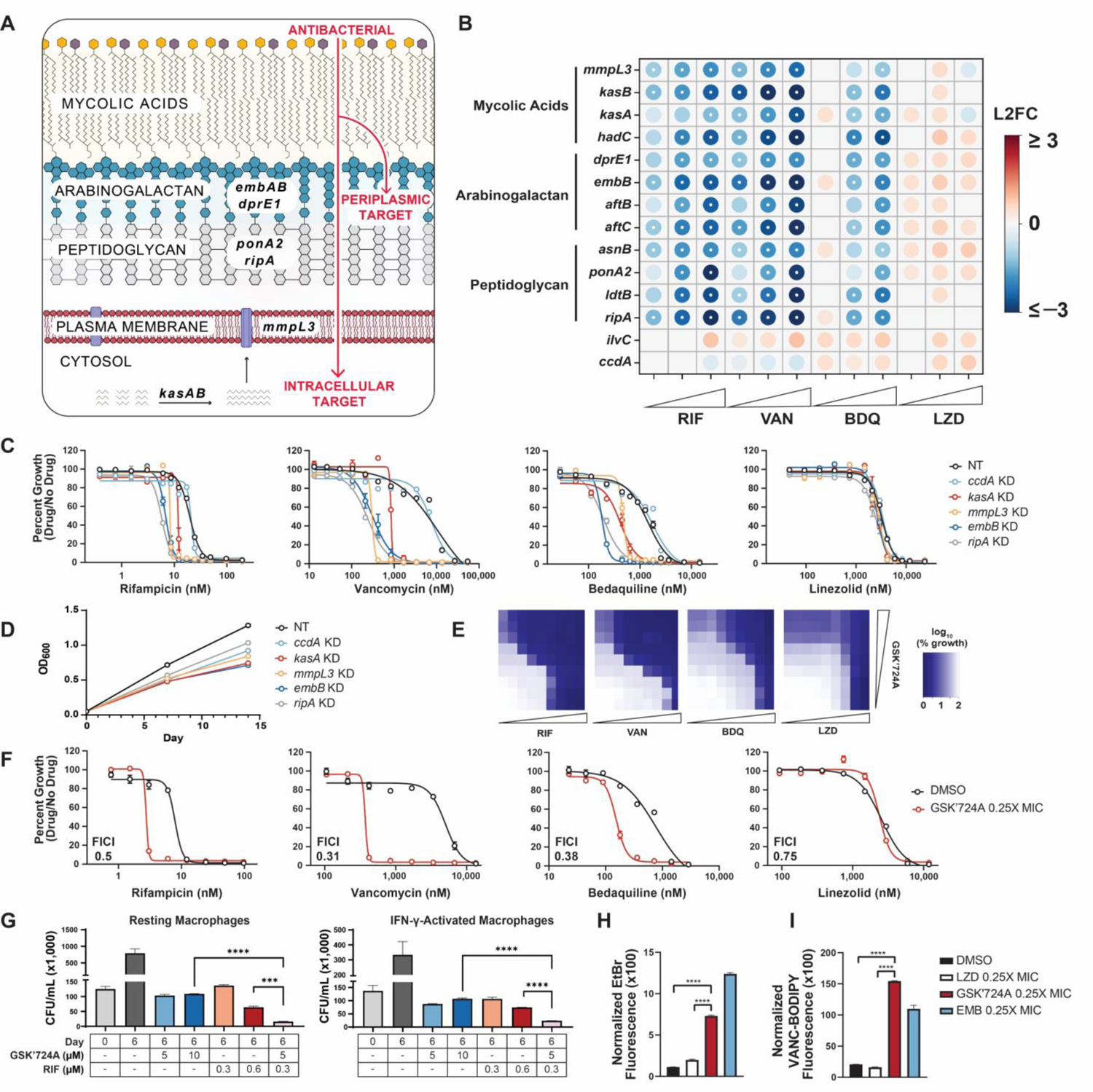
The Mtb envelope mediates intrinsic resistance to a subset of drugs. (A) Diagram of the mycobacterial mAGP complex. Select genes involved in mycolic acid synthesis and transport (*kasAB, mmpL3*), arabinogalactan biosynthesis (*embAB, dprE1*), and peptidoglycan remodeling (*ponA2, ripA*) are highlighted. (B) Feature-expression heatmap of select chemical-genetic hit genes for the indicated drugs from the 5-day CRISPRi library pre-depletion screen. The color of each circle represents the gene-level L2FC. A white dot represents an FDR < 0.01 and a |L2FC| > 1. *ilvC* and *ccdA* are included as non-hit controls. (C-D) Single strain validation of mAGP-associated hits. MIC values (C) for the indicated drugs were measured for hypomorphic CRISPRi strains targeting *kasA*, *mmpL3*, *embB*, *ripA*, and the non-hit essential gene *ccdA*. Growth curves (D) are derived from the vehicle control samples. NT corresponds to a CRISPRi strain harboring a non-targeting sgRNA. Data represent mean ± SEM for technical triplicates. Data are representative of at least two independent experiments. KD = knockdown. (E-F) KasA inhibitor (GSK’724A) checkerboard assays to quantify drug-drug interactions. MIC curves are shown for each drug in the absence (DMSO) or presence of 0.25X MIC_80_ GSK’724A. The fractional inhibitory concentration index (FICI) values listed represent the lowest value obtained from each checkerboard assay. Error bars represent the SEM for technical triplicates. Data are representative of at least two independent experiments. (G) GSK’724A synergy with rifampicin in resting and IFN-γ-activated murine bone marrow derived macrophages. Mtb-infected macrophages were treated with the indicated concentrations of GSK’724A or rifampicin for 6 days prior to plating for colony forming units (CFU). Data represent mean ± SEM for technical triplicates. Results from an unpaired t-test are shown: ***, p < 0.001, ****, p < 0.0001. Data are representative of two independent experiments. (H-I) Ethidium bromide (H) and Vancomycin-BODIPY (I) uptake of H37Rv pre-treated for two days with DMSO or subinhibitory linezolid, GSK’724A, or ethambutol. Data represent mean ± SEM for 4 replicates. Results from an unpaired t-test are shown: ****, p<0.0001.

**Supplemental Figure 5:**
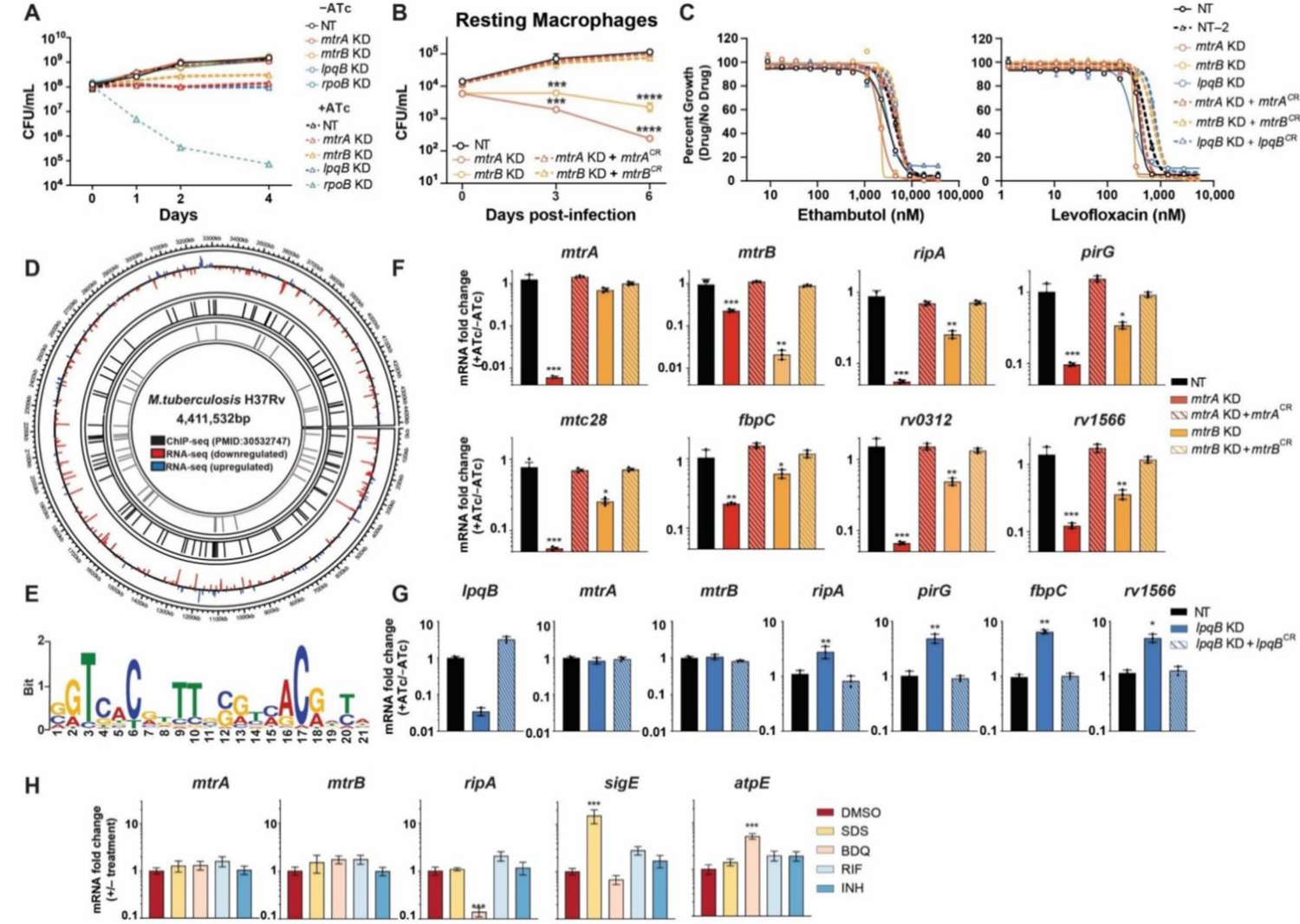
The MtrAB two-component system is critical for multi-drug intrinsic resistance in Mtb. (A) Time-kill curves for the indicated CRISPRi strains. Data represent mean ± SEM for technical triplicates. Data are representative of at least two independent experiments. NT = non-targeting; KD = knockdown; CR = CRISPRi-resistant. (B) Growth of the indicated CRISPRi strains in resting murine bone marrow derived macrophages. Bacterial strains were exposed to ATc (100 ng/mL) for 24 hours prior to macrophage infection. 3 and 6 days after infection, bacteria were harvested and quantified by CFU. Data represent mean ± SEM for technical triplicates. Significance was determined by two-way ANOVA and adjusted for multiple comparisons. ***, p<0.001; ****, p<0.0001. (C) MIC values for the indicated drugs were measured against the indicated strains. Data represent mean ± SEM for technical triplicates. Data are representative of at least two independent experiments. (D) Circos plot depicting overlapping genes identified by RNA-seq (**Figure 2F**) and MtrA ChIP-seq (Gorla et al., 2018). Outer track: the H37Rv genome by nucleotide position; middle track: lines mark genes with a significant L2FC values (padj< 0.05) upon *mtrA* knockdown (blue = positive L2FC; red = negative L2FC); inner tracks: black lines mark genes defined as interacting with MtrA by ChIP-seq (Gorla et al., 2018), and grey lines highlight genes which display both a significant L2FC (padj< 0.05; |L2FC| > 1) by *mtrA* RNAseq and are found to interact with MtrA by ChIP-seq. (E) Identification of an MtrA consensus binding motif. MEME analysis (Bailey et al., 2009) was performed on the promoter regions of candidate genes found to both be downregulated upon *mtrA* silencing (**Figure 2F**) and bound by MtrA by ChIP-seq (Gorla et al., 2018) (n=25 genes). (F,G) Quantification of indicated gene mRNA levels by qRT-PCR. Strains were grown in the presence of ATc for ∼3 generations prior to harvesting RNA. Error bars are SEM of three technical replicates. Statistical significance was calculated as p-value with unpaired T-test. *, p<0.05; **, p<0.01; ***, p<0.001. (H) Quantification of indicated gene mRNA levels by qRT-PCR. Wild-type H37Rv was grown in the presence of the indicated stress (RIF/BDQ/INH: 4 x IC_50_, SDS: 0.2%, DMSO: 0.5%) for 3 hours prior to harvesting RNA. Error bars are SEM of three technical replicates. Statistical significance was calculated as p-value with unpaired T-test. ***, p<0.001.

**Supplemental Figure 6:**
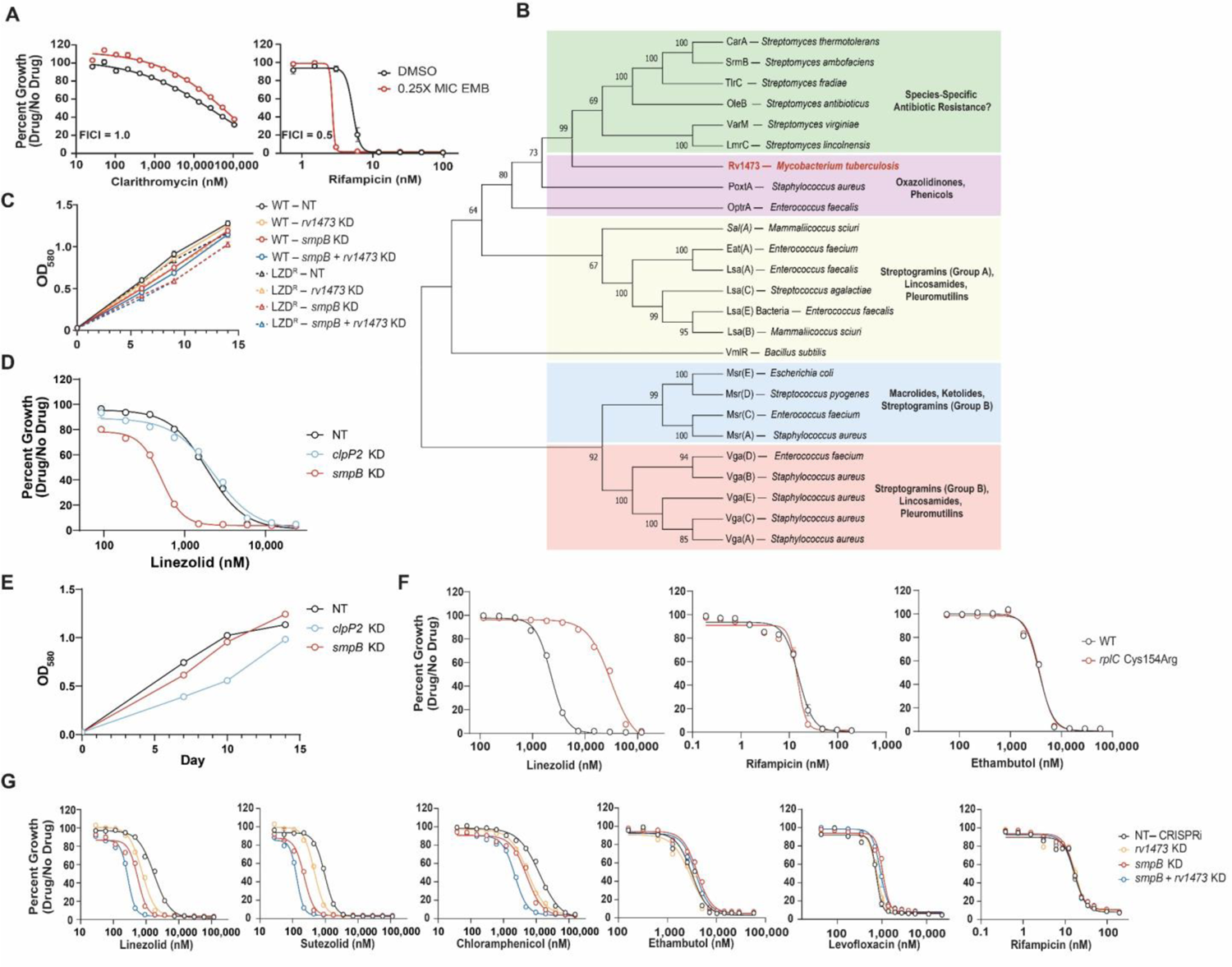
Mtb encodes diverse mechanisms of intrinsic resistance to ribosome-targeting antibiotics. (A) Ethambutol checkerboard assays to quantify drug-drug interactions. MIC curves are shown for each drug in the absence (DMSO) or presence of 0.25X MIC_80_ of EMB. Fractional inhibitory concentration index (FICI) values listed represent the lowest value obtained from each checkerboard assay. Data represent mean ± SEM for technical triplicates. (B) Phylogenetic tree of antibiotic resistance (ARE) ABC-F proteins from the indicated species. Figure adapted from (Sharkey et al., 2016). Bootstrap values (500 replicates) are indicated at each node. (C) Growth curves for the LZD-associated hit genes and control strains shown in **Figure 3D**. Curves are derived from the vehicle control samples of the MIC assay. Data represent mean ± SEM for technical triplicates. Results are representative data from at least two independent experiments. NT = non-targeting; KD = knockdown. (D) MIC values for LZD were measured for CRISPRi knockdown strains targeting *smpB* and *clpP2* in wild-type H37Rv. Data represent mean ± SEM for technical triplicates. (E) Growth curves for the strains shown in (D). Curves are derived from the vehicle control samples of the MIC assay. Data represent mean ± SEM for technical triplicates. (F) MIC curves of WT H37Rv and an isogenic *rplC*-Cys154Arg mutant for LZD, RIF, and EMB. Data represent mean ± SEM for six technical replicates. (G) MIC values for the indicated drugs were measured for the indicated CRISPRi strains. Data represent mean ± SEM for technical triplicates.

**Supplemental Figure 7:**
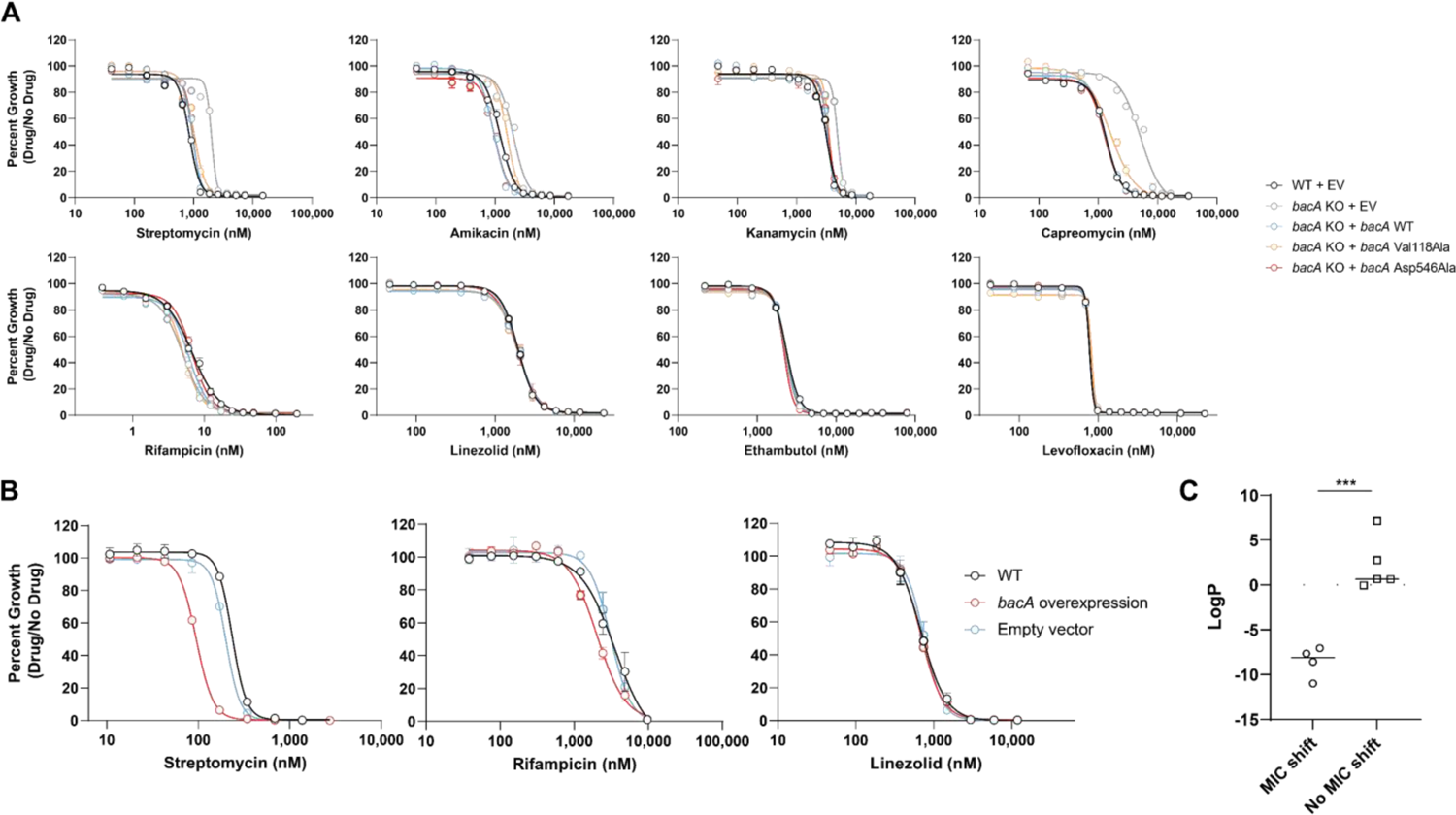
Loss-of-function mutations in *bacA* confer resistance to aminoglycosides and capreomycin. (A) MIC values for the indicated drugs were measured for the indicated strains. Data represent mean ± SEM for technical triplicates. Results are representative data from at least two independent experiments. (B) Overexpression of Mtb *bacA* confers streptomycin sensitivity in *M. smegmatis*. MIC values for the indicated drugs were measured for the three indicated strains. Data represent mean ± SEM for technical triplicates. (C) LogP values for the antibiotics to which *bacA* mutants show an increased MIC (STR, AMK, CAP, KAN) or no MIC change (RIF, EMB, LVX, LZD). Results from an unpaired t-test are shown: ***, p<0.001.

**Supplemental Figure 8:**
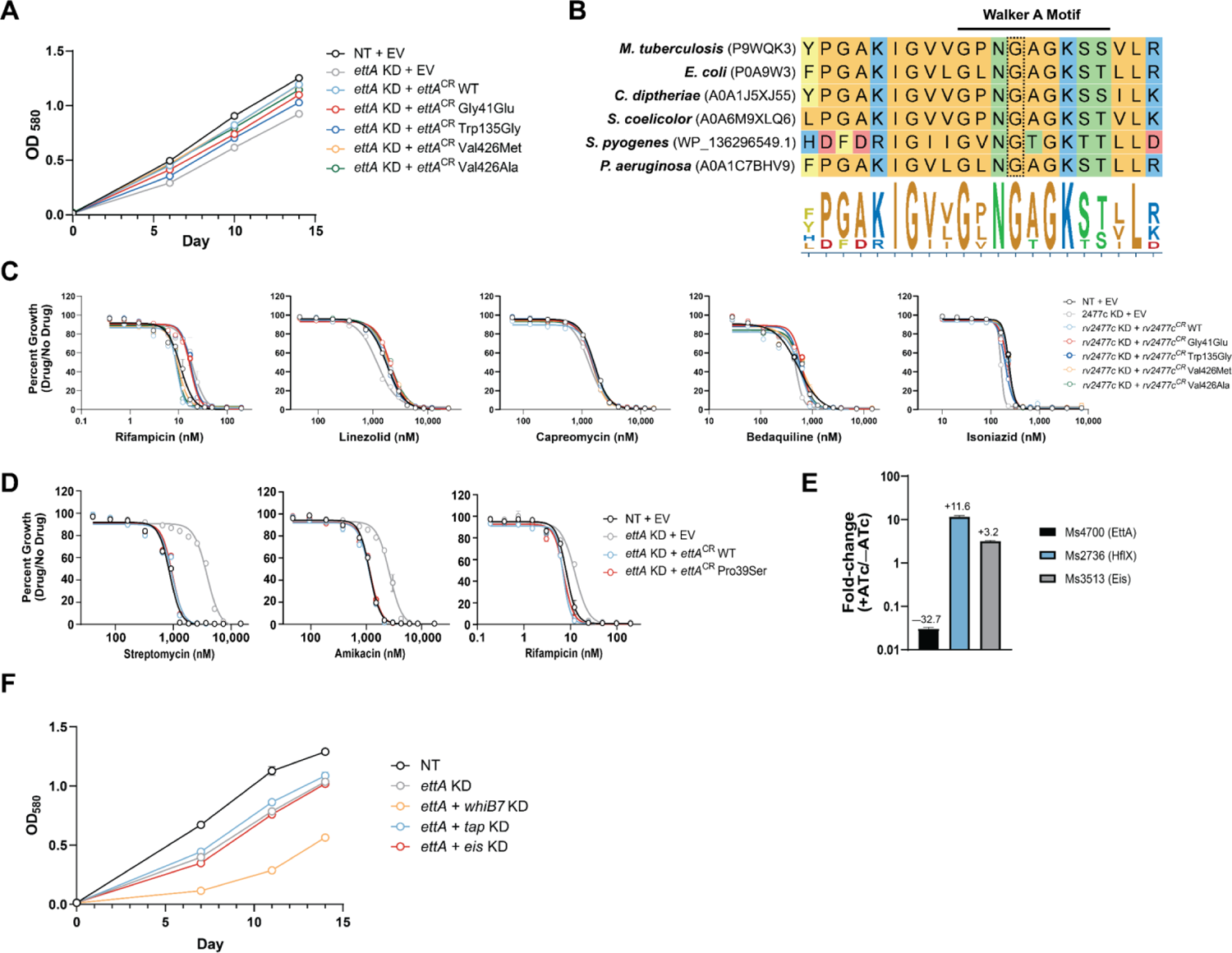
Partial loss-of-function mutations in *ettA* confer low-level multidrug resistance and are associated with an MDR outbreak in South America. (A) Growth curves for the strains shown in **Figure 5C**. Curves are derived from the vehicle control samples of the MIC assay. Data represent mean ± SEM for technical triplicates. Results are representative data from at least two independent experiments. (B) Amino acid alignment for EttA orthologs from the indicated species for the region surrounding the N-terminal Walker A motif. The Mtb EttA Gly41 residue is boxed. Accession numbers are listed next to each species. (C) MIC values for the indicated drugs were measured for the strains shown in **Figure 5C**. Data represent mean ± SEM for technical triplicates. Results are representative data from at least two independent experiments. (D) The Pro39Ser mutation in *ettA* does not confer antibiotic resistance. MIC values for the indicated drugs were measured as in **Figure 5C**. Data represent mean ± SEM for technical triplicates. (E) Quantitative mass spectrometry results from experiments described in (Bosch et al., 2021). Values indicate protein level fold-change following CRISPRi knockdown of *ms4700*. Data represent mean ± SEM four technical replicates derived from two biological replicates. Ms4700 could only be detected in two replicates and, thus, the mean ± SEM for duplicates is shown. Growth curves for the strains shown in **Figure 5E**. Curves are derived from the vehicle control samples of the MIC assay. Data represent the mean ± SEM for technical triplicates.

**Supplemental Figure 9:**
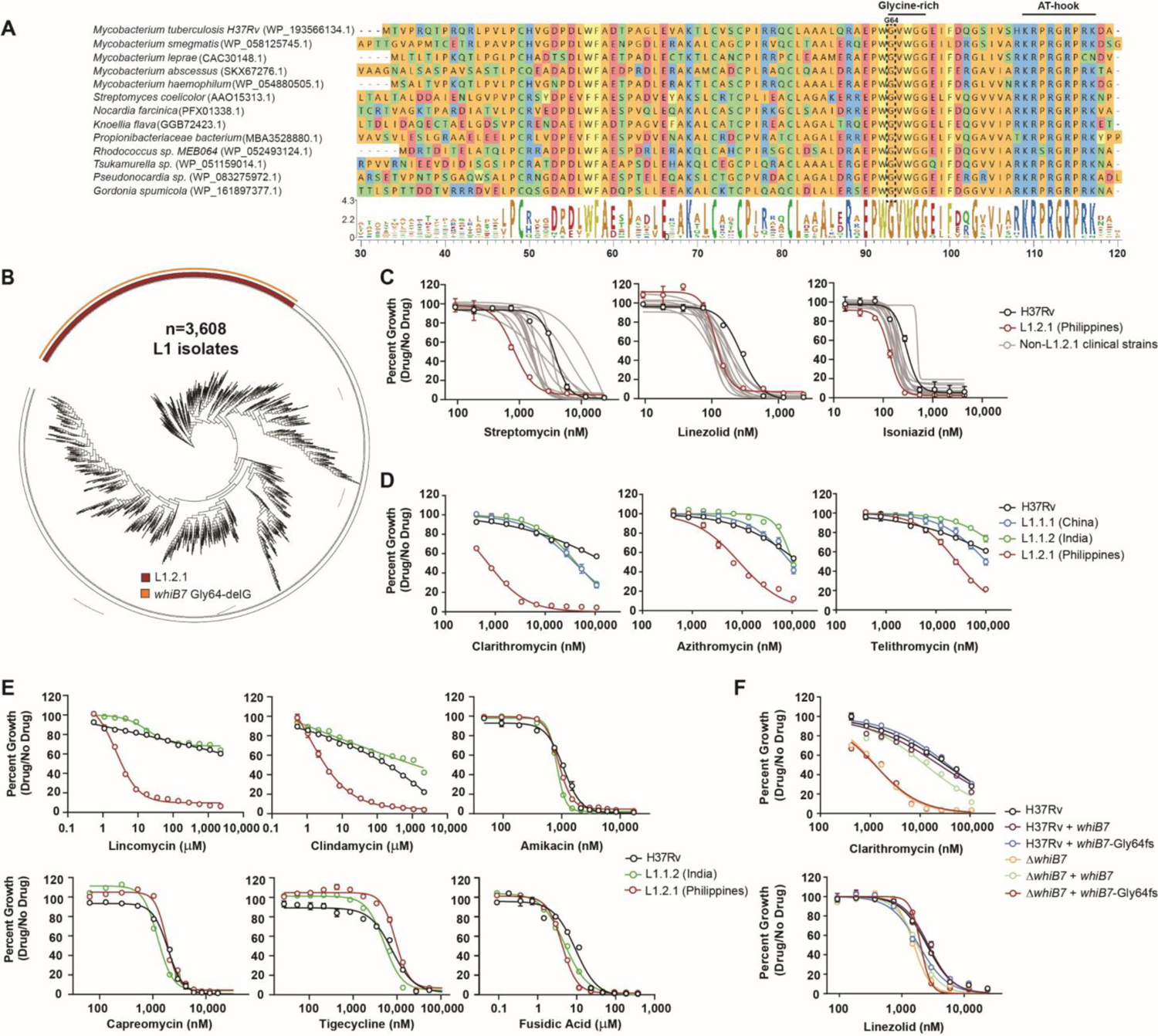
The L1.2.1 sub-lineage has a loss-of-function mutation in *whiB7* that renders it hypersusceptible to macrolides, ketolides, and lincosamides. (A) Alignment of WhiB7 orthologues from representative actinobacteria. Accession numbers are listed next to each species. The conserved glycine-rich motif and DNA binding AT-hook element are highlighted. (B) Phylogenetic tree of all L1 Mtb clinical isolates (n=3,608) in our WGS database (**Supplemental Data 4**). L1.2.1 and the *whiB7* Gly64delG mutation are highlighted. (C-E) MIC values the indicated drugs were measured for a reference set of Mtb clinical strains. Error bars represent the standard error of the mean (SEM) for technical triplicates. Results are representative data from at least two independent experiments. (F) H37Rv was transformed with an integrating plasmid to express either the H37Rv *whiB7* allele or the L1.2.1 *whiB7* Gly64-delG allele *in trans* from its native promoter. The resulting strains were then used to measure MIC values for the indicated drugs. Error bars represent the SEM for technical triplicates. Results are representative data from at least two independent experiments.

**Supplemental Figure 10:**
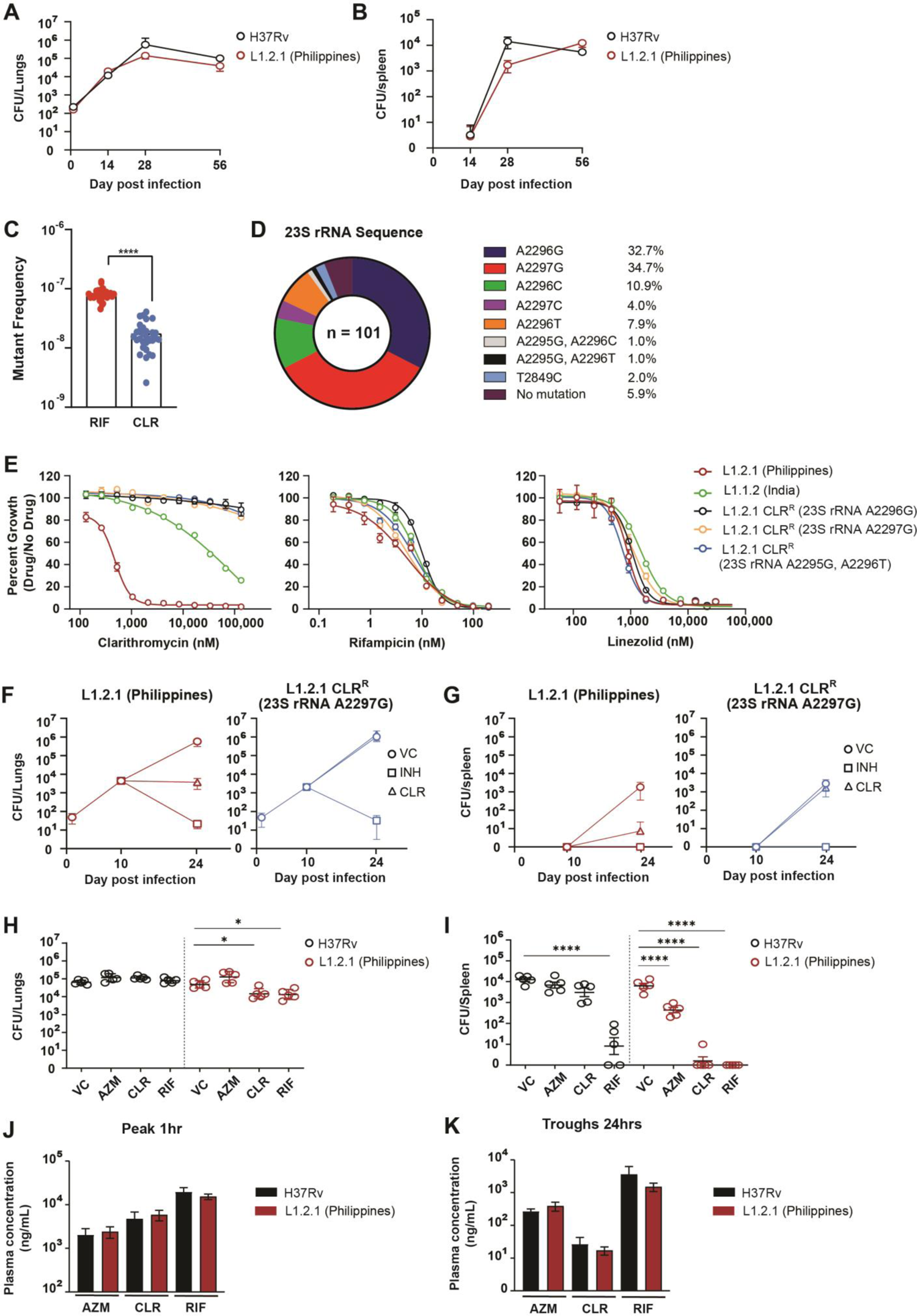
The L1.2.1 sublineage is susceptible to clarithromycin *in vivo*. (A,B) Growth kinetics of H37Rv and L1.2.1 *in vivo*. BALB/c mice were infected with approximately 100-200 CFU by aerosol and killed over the course of infection at indicated time points. Mean lung (A) and spleen (B) Mtb CFU (± SEM) in BALB/c mice were determined after primary infection (C) Rifampicin and clarithromycin mutation frequency analysis with the L1.2.1 strain. (D) Distribution of 23S rRNA mutations from *in vitro*-selected, clarithromycin-resistant L1.2.1 isolates from panel (C). (E) MIC profiles of representative CLR-resistant L1.2.1 isolates. Error bars represent the standard error of the mean (SEM) for technical triplicates. Results are representative data from at least two independent experiments. (F,G) Mean lung (F) and spleen (G) Mtb CFU (± SEM) in BALB/c mice after isoniazid (INH; 25 mg/kg), or clarithromycin (CLR; 200 mg/kg) treatment. Mice were infected with approximately 100-200 CFU of aerosolized Mtb. After ten days to allow the acute infection to establish, chemotherapy was initiated. Mtb bacterial load of lungs and spleen were determined at the indicated time points. VC = vehicle control. CLR^R^ = clarithromycin-resistant. (H,I) Mean lung (F) and spleen (G) Mtb colony-forming units (CFU; ± SEM) in BALB/c mice after azithromycin (AZM; 200 mg/kg), clarithromycin (CLR; 200 mg/kg), or rifampicin (RIF; 25 mg/kg) treatment. Mice were infected with approximately 100-200 CFU of aerosolized Mtb. After two weeks to allow the acute infection to establish, chemotherapy was initiated. Following two weeks of drug therapy, Mtb bacterial load of lungs and spleen were determined. Statistical significance was assessed by one-way ANOVA followed by Tukey’s post-hoc test. *, p <0.05; ****, p <0.0001. (J,K) Monitoring of plasma drug concentrations after 2 weeks of therapy, prior to CFU enumeration in lungs and spleen. Blood was collected at 1h (J) and 24h (K) post-dose after 13 daily doses, from 4 mice in each infection and treatment group described in (H,I). Drug concentrations were measured in plasma using high pressure liquid chromatography coupled to tandem mass spectrometry. Mean and standard deviation (error bars) are shown.

**Supplemental Figure 11:**
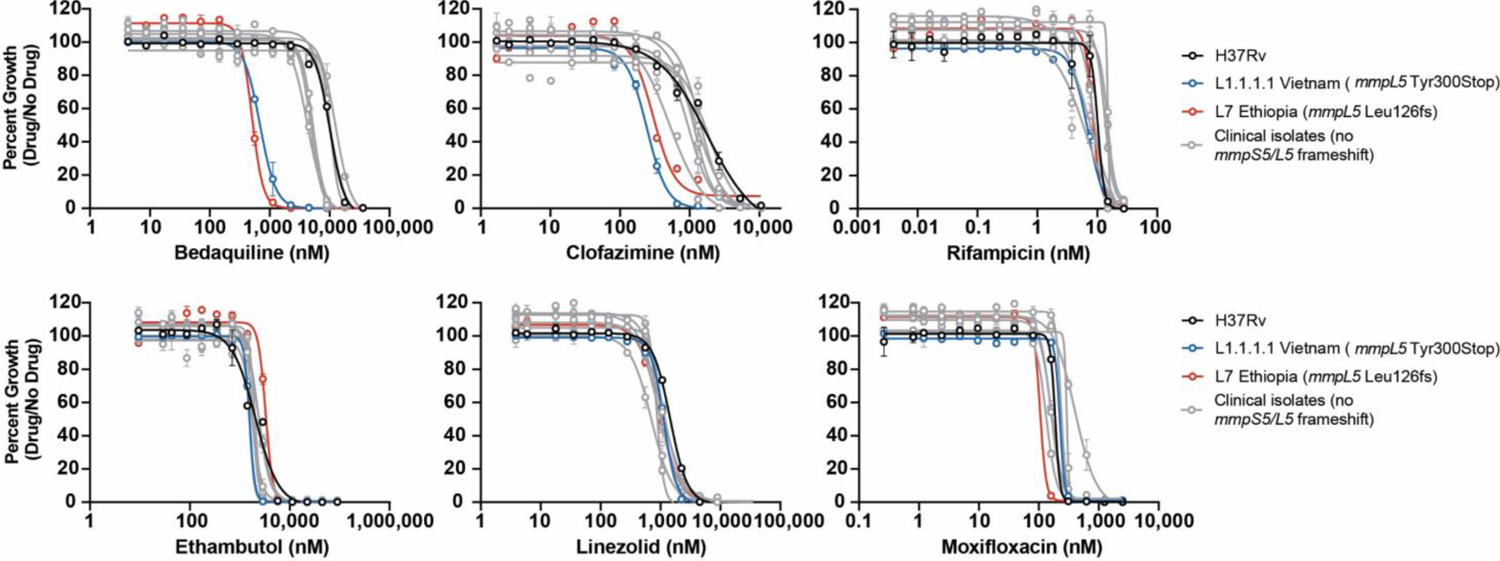
Loss-of-function mutations on MmpL5 renders it sensitive to Bedaquiline and Clofazimine MIC values the indicated drugs were measured for a reference set of Mtb clinical strains. Error bars represent the standard error of the mean (SEM) for technical triplicates. Results are representative data from at least two independent experiments.

## SUPPLEMENTAL INFORMATION

**Supplemental Table 1:**
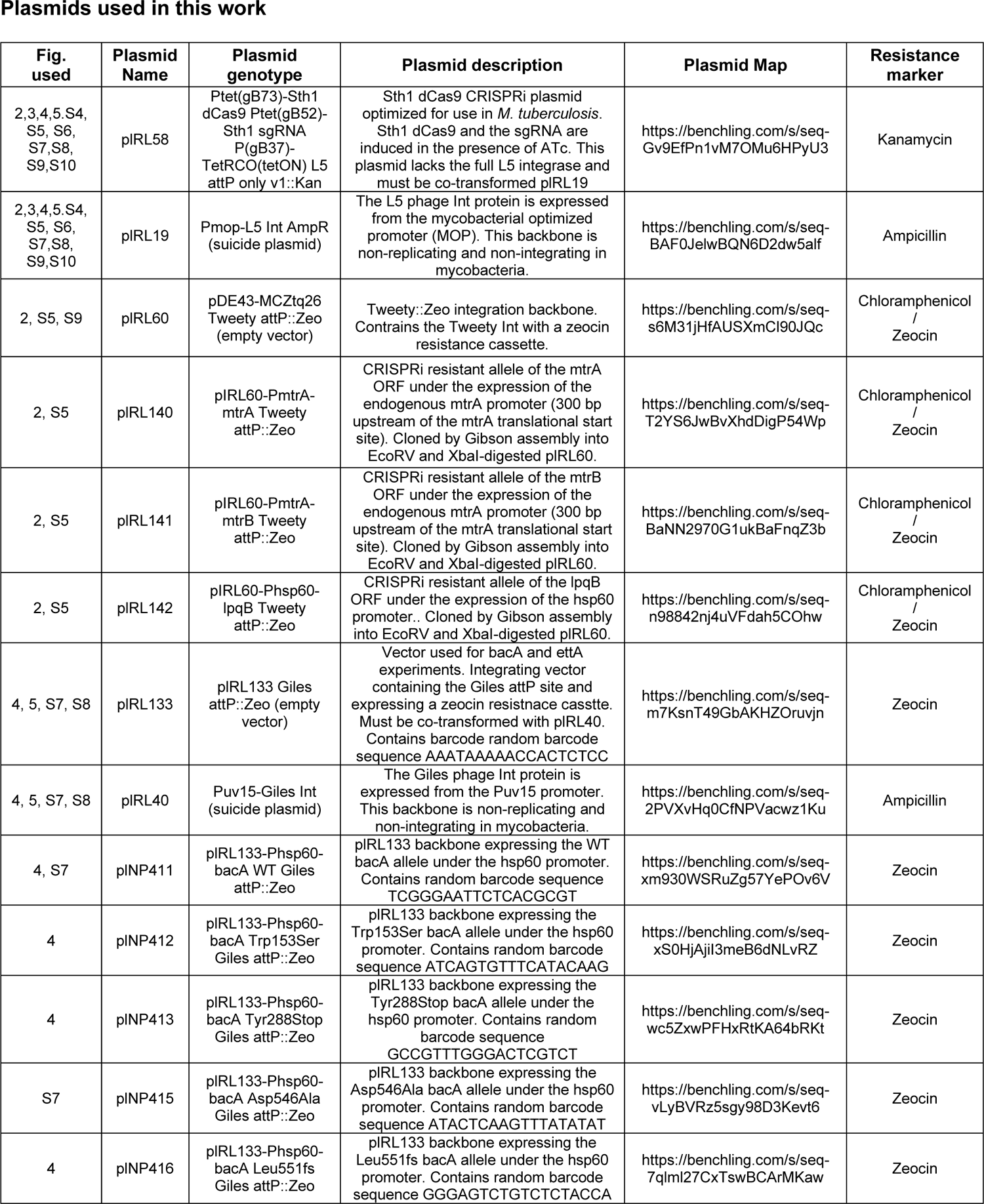
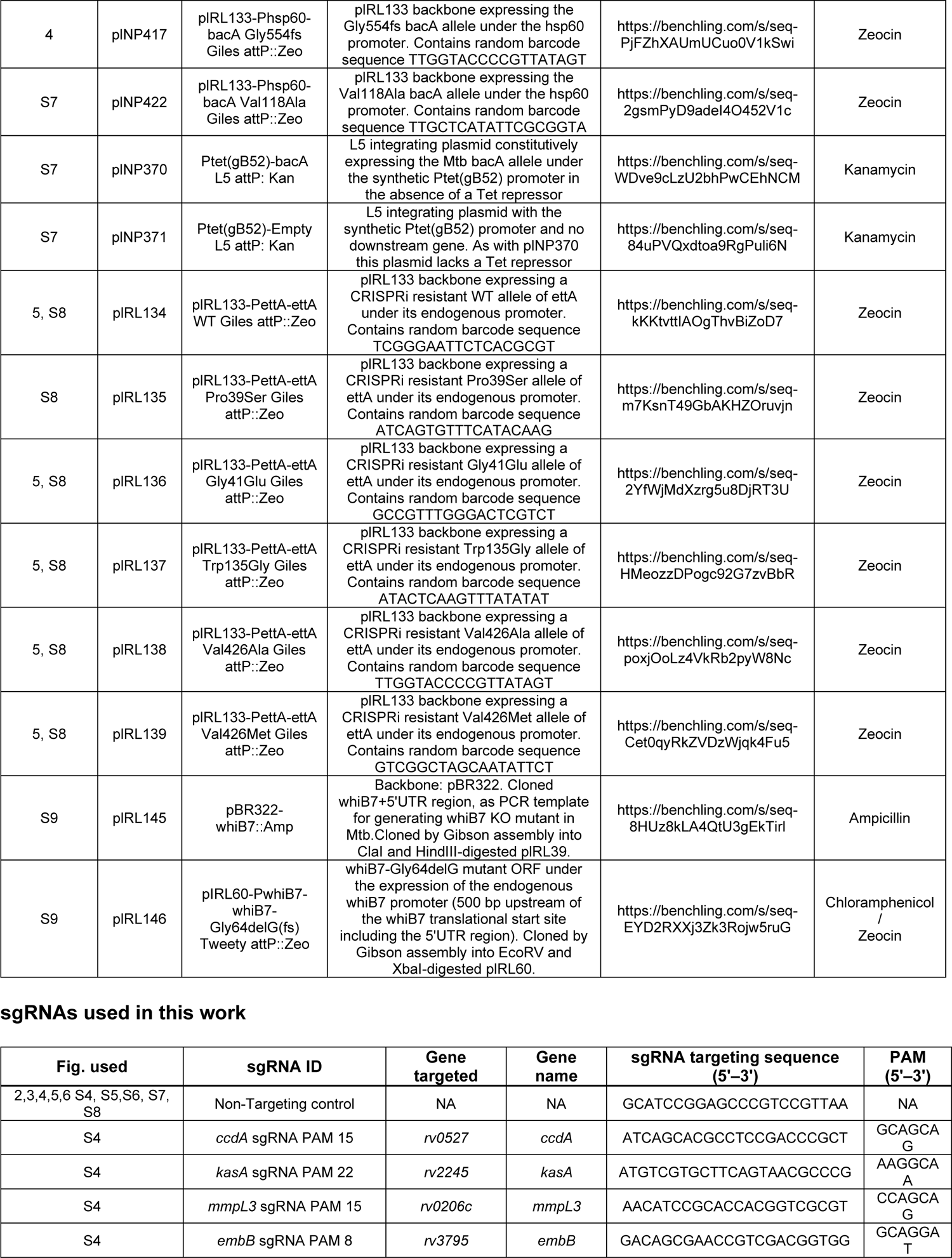
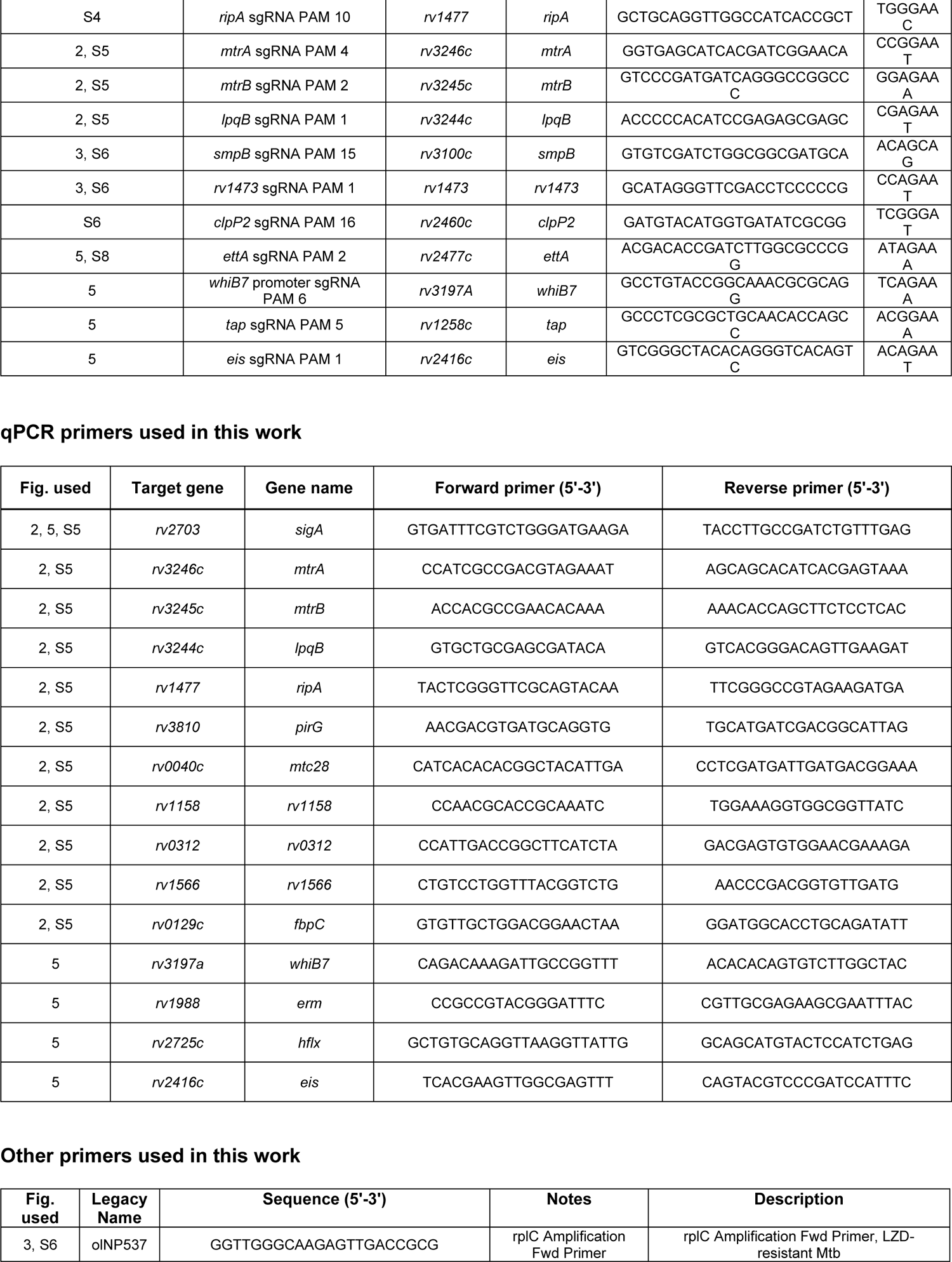
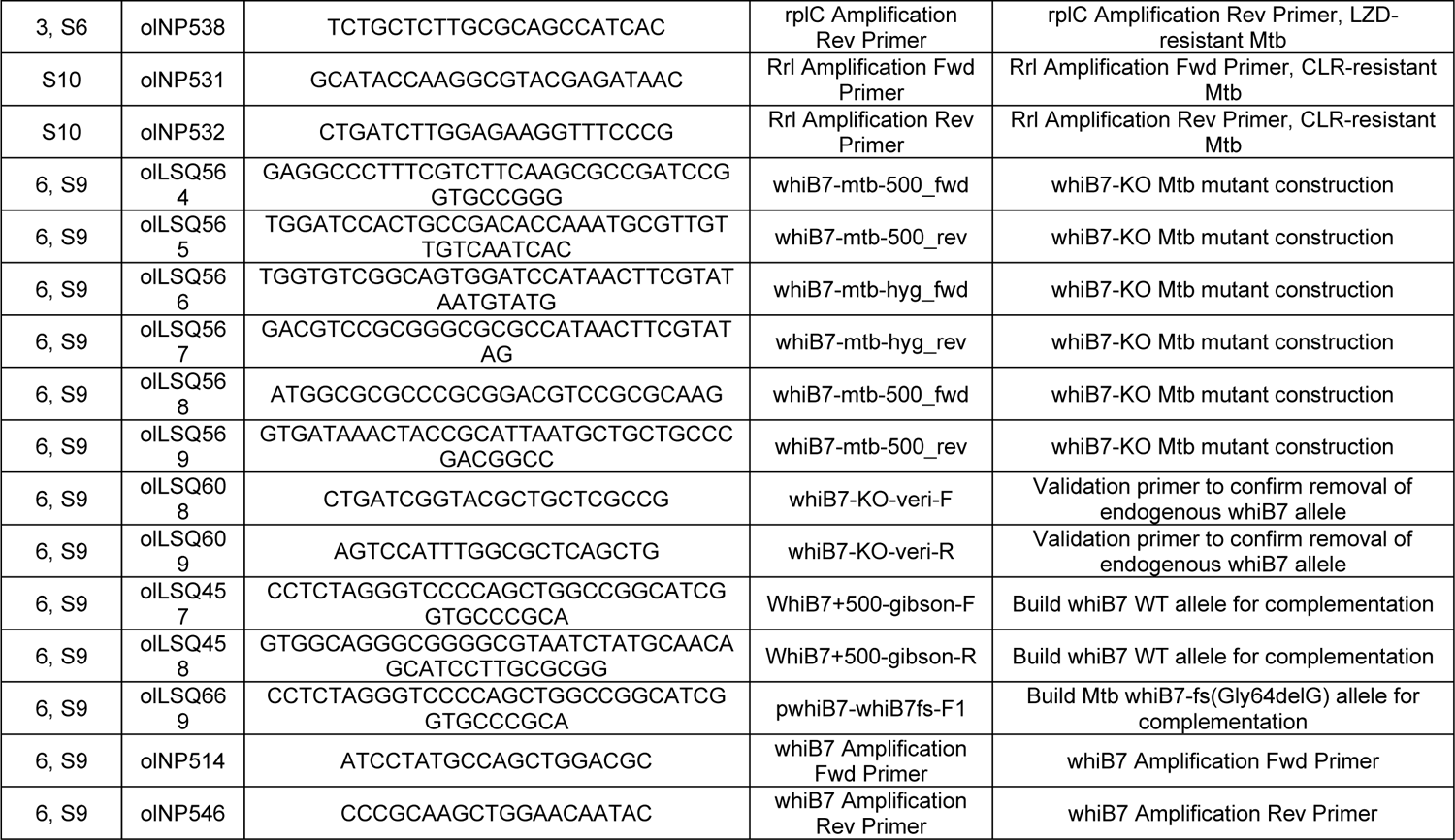
List of plasmids and primers used in this work

**Supplemental Table 2:** Additional acquired drug sensitivity candidates

| Gene | Drug | Mutation | Sublineage | Occurrences | Notes |
| --- | --- | --- | --- | --- | --- |
| *whiB7* | CLR | Leu18fs | lineage2.2.1: 100/6719 | 100 |  |
| *whiB7* | CLR | Ala60fs | lineage4: 2/814, lineage4.8: 18/3234 | 20 |  |
| *hflX* | CLR | Glu206fs | lineage4.9: 93/2941 | 93 |  |
| *hflX* | CLR | Arg237fs | lineage4.9: 12/2941 | 12 |  |
| *hflX* | CLR | Gln367Stop | lineage4.2.1: 117/666, lineage4.6: 2/187 | 119 |  |
| *rv2369c* | CLR | Arg57fs | None: 12/3241 | 12 | Possible polar effect with *phoH1* |
| *rv2369c* | CLR | Cys58fs | lineage3: 11/3537 | 11 | Possible polar effect with *phoH1* |
| *rv2369c* | CLR | Gln72Stop | lineage1.2.1: 2/164, lineage1.2.1.3: 78/435 | 80 | Possible polar effect with *phoH1* |
| *mm pL5* | CLR/<br>BDQ | Asp132fs | None: 48/3241, lineage1.2.1: 1/164, lineage2.2.1: 5/6719, lineage4.1.1.3: 2/1178, lineage4.8: 1/3166 | 57 | Also observed by Merker et al., may also confer clofazamine sensitivity(Merker et al., 2020) |
| *mm pL5* | CLR/<br>BDQ | Arg202fs | lineage4: 55/807 | 55 | Also observed by Merker et al., may also confer clofazamine sensitivity(Merker et al., 2020) |
| *mm pL5* | CLR/<br>BDQ | Tyr300Stop | lineage1.1.1: 4/54, lineage1.1.1.1: 75/108, lineage4.8: 1/3166 | 80 | Also observed by Merker et al., may also confer clofazamine sensitivity(Merker et al., 2020) |
| *mm pL5* | CLR/<br>BDQ | Pro498fs | lineage2.2.2: 1/747, lineage4.6: 59/187 | 60 | Also observed by Merker et al., may also confer clofazamine sensitivity(Merker et al., 2020) |
| *rv3822* | BDQ | Leu123fs | None: 40/2223 | 40 |  |
| *rv3822* | BDQ | Tyr208fs | lineage4: 4/814, lineage4.8: 36/3234 | 40 |  |

**Supplemental Data 1:** MAGeCK screen results

**Supplemental Data 2:** Chemical-genetic hit overlap between CRISPRi and TnSeq

**Supplemental Data 3:** RNA-seq differential expression analysis results for Mtb *mtrA* CRISPRi knockdown mutant

**Supplemental Data 4**: Full list of NCBI accession numbers of WGS sequences used in this study

**Supplemental Data 5**: List of identified *tsnR* mutations from clinical isolate WGS database and co-occurrence with *rplC* Cys154Arg Linezolid-resistant isolates

**Supplemental Data 6**: List of identified *bacA* mutations from clinical isolate WGS database

**Supplemental Data 7**: List of identified *ettA* mutations from clinical isolate WGS database

**Supplemental Data 8**: List of identified *whiB7* mutations from clinical isolate WGS database

**Supplemental Data 9**: Acquired drug resistance candidate mutations

## MATERIALS AND METHODS

### Bacterial strains

Mtb strains are derivatives of H37Rv unless otherwise noted. A reference set of Mtb clinical strains was obtained from the Belgian Coordinated Collections of Microorganisms (BCCM) (Borrell et al., 2019). *E. coli* strains are derivatives of DH5alpha (NEB).

### Mycobacterial cultures

Mtb was grown at 37°C in Difco Middlebrook 7H9 broth or on 7H10 agar supplemented with 0.2% glycerol (7H9) or 0.5% glycerol (7H10), 0.05% Tween-80, 1X oleic acid-albumin-dextrose-catalase (OADC) and the appropriate antibiotics. Where required, antibiotics or small molecules were used at the following concentrations: kanamycin at 20 μg/mL; anhydrotetracycline (ATc) at 100 ng/mL, and zeocin at 20 μg/mL. Mtb cultures were grown standing in tissue culture flasks (unless otherwise indicated) at 37°C, 5% CO_2_.

### Generation of individual CRISPRi and CRISPRi-resistant complementation strains

Individual CRISPRi plasmids were cloned as previously described in (Bosch et al., 2021) using Addgene plasmid #166886. Briefly, the CRISPRi plasmid backbone was digested with BsmBI-v2 (NEB #R0739L) and gel purified. sgRNAs were designed to target the non-template strand of the target gene ORF. For each individual sgRNA, two complementary oligonucleotides with appropriate sticky end overhangs were annealed and ligated (T4 ligase NEB # M0202M) into the BsmBI-digested plasmid backbone. Successful cloning was confirmed by Sanger sequencing.

Individual CRISPRi plasmids were then electroporated into Mtb. Electrocompetent cells were obtained as described in(Murphy et al., 2015). Briefly, a WT Mtb culture was expanded to an OD_600_=0.8-1.0 and pelleted (4,000 x g for 10 min). The cell pellet was washed three times in sterile 10% glycerol. The washed bacilli were then resuspended in 10% glycerol in a final volume of 5% of the original culture volume. For each transformation, 100 ng plasmid DNA and 100 μL of electrocompetent mycobacteria were mixed and transferred to a 2 mm electroporation cuvette (Bio-Rad #1652082). Where necessary, 100 ng of plasmid plRL19 (Addgene plasmid #163634) was also added. Electroporation was performed using the Gene Pulser X cell electroporation system (Bio-Rad #1652660) set at 2500 V, 700 Ω and 25 μF. Bacteria were recovered in 7H9 for 24 hours. After the recovery incubation, cells were plated on 7H10 agar supplemented with the appropriate antibiotic to select for transformants.

To complement CRISPRi-mediated gene knockdown, synonymous mutations were introduced into the complementing allele at both the protospacer adjacent motif (PAM) and seed sequence (the 8-10 most PAM-proximal bases at the 3’ end of the sgRNA targeting sequence) to prevent sgRNA targeting. Silent mutations were introduced into Gibson assembly oligoes to generate these “CRISPRi resistant” (CR) alleles. Complementation alleles were expressed from the endogenous or hsp60 promoters in a Tweety or Giles integrating plasmid backbone, as indicated in each figure legend and/or the relevant plasmid maps (**Supplemental Table 1**). These alleles were then transformed into the corresponding CRISPRi knockdown strain, with the plRL40 Giles Int expressing plasmid where necessary.

The full list of sgRNA targeting sequences and complementation plasmids can be found in **Supplemental Table 1**.

### Construction of the Δ*whiB7* and complemented Mtb strains

The Mtb Δ*whiB7* strain was constructed by allelic exchange using a RecET-mediated recombineering approach as previously described(Murphy et al., 2015). Deletion of *whiB7* was confirmed by PCR and whole-genome sequencing (BGI). The Δ*whiB7* strain was complemented by reintroducing a wild-type copy of *whiB7* under the control of its native promoter at the *attL5* site of the Mtb genome. Plasmid sequences and maps can be found in **Supplemental Table 1**.

### Pooled CRISPRi chemical-genetic screening

Chemical-genetic screens were initiated by thawing 5 × 1 mL (1 OD_600_ unit per mL) aliquots of the Mtb CRISPRi library (RLC12; Addgene #163954) and inoculating each aliquot into 19 mL 7H9 supplemented with kanamycin (10 μg/ml) in a vented tissue culture flask flask (T-75; Falcon #353136). The starting OD_600_ of each culture was approximately 0.05. Cultures were expanded to OD_600_=1.5, pooled and passed through a 10 μm cell strainer (pluriSelect #43-50010-03) to obtain a single cell suspension. The single cell suspension was then treated with ATc (100 ng/mL final concentration) to initiate target pre-depletion. To generate 1 day pre-depletion culture, the single-cell suspension was diluted back to OD_600_=0.5 in a total volume of 40 mL 7H9. The remaining single-cell suspension was used to generate a 5-day pre-depletion culture, with a starting OD_600_=0.1 (40 mL; 100 ng/mL ATc). After 4 days, the 5-day pre-depletion start culture was further diluted back to a starting OD_600_=0.05 (40 mL; in 100 ng/mL ATc) and incubated for a further 5 days to generate the 10 day pre-depletion culture.

To initiate the chemical-genetic screen, we first harvested 10 OD_600_ units of bacteria (∼3×10^9^ bacteria; ∼30,000X coverage of the CRISPRi library) from the 1, 5, or 10-day CRISPRi library pre-depletion cultures as input controls. Triplicate cultures were then inoculated at OD_600_=0.05 in 10 ml 7H9 supplemented with ATc (100 ng/mL), kanamycin (10 μg/ml), and the indicated drug concentration or DMSO vehicle control (see **Supplemental Figure 1**). Pooled CRISPRi chemical-genetic screens were performed in vented tissue culture flasks (T-25; Falcon #353109). Cultures were outgrown for 14 days at 37°C, 5% CO_2_. ATc was replenished at 100 ng/mL at day 7. After 14 days outgrowth, OD_600_ values were measured for all cultures to empirically determine the MIC for each drug. Samples from three descending doses of partially inhibitory drug concentrations were processed for genomic DNA extraction, defined as “High”, “Medium”, and “Low” in **Supplemental Figure 1**, as described below. Due to an error during genomic DNA extraction, EMB 1,804 nM (“Med”) Day 1 data reflects two biological replicates, one of which was sequenced twice to produce 3 replicates. Additionally, the 10-day sample was lost for the 221 nM (“Med”) streptomycin screen.

### Genomic DNA extraction and library preparation for Illumina sequencing

Genomic DNA was isolated from bacterial pellets using the CTAB-lysozyme method as previously described(Bosch et al., 2021). Briefly, Mtb pellets (5-30 OD_600_ units) were resuspended in 1 mL of PBS + 0.05% Tween-80. Cell suspensions were centrifuged for 5 min at 4,000 x g and the supernatant was removed. Pellets were resuspended in 800 µL TE buffer (10 mM Tris pH 8.0, 1 mM EDTA) + 15 mg/mL lysozyme (Alfa Aesar J60701-06) and incubated at 37°C for 16 hours. Next, 70 µL of 10% SDS (Promega V6551) and 5 µL of proteinase K (20 mg/mL, Thermo Fisher 25530049) were added and samples were incubated at 65°C for 30 min. Subsequently, 100 µL of 5 M NaCl and 80 µL of 10% CTAB (Sigma Aldrich H5882) were added and samples were incubated for an additional 30 min at 65°C. Finally, 750 µL of ice-cold chloroform was added and samples were mixed. After centrifugation at 16,100 x g and extraction of the aqueous phase, samples were removed from the biosafety level 3 facility. Samples were then treated with 25 µg of RNase A (Bio Basic RB0474) for 30 min at 37°C followed by an extraction with phenol:chloroform:isoamyl alcohol (pH=8.0, 25:24:1 Thermo Fisher BP1752I-400) then chloroform.

Genomic DNA was precipitated from the final aqueous layer (600 µL) with the addition of 10 µL of 3 M sodium acetate and 360 µL of isopropanol. DNA pellets were spun at 21,300 x g for 30 min at 4°C and washed 2X with 750 µL of 80% ethanol. Pellets were dried and resuspended with elution buffer (Qiagen 19086) before spectrophotometric quantification.

The concentration of isolated genomic DNA was quantified using the DeNovix dsDNA high sensitivity assay (KIT-DSDNA-HIGH-2; DS-11 Series Spectrophotometer / Fluorometer). Next, the sgRNA-encoding region was amplified from 500 ng genomic DNA with 17 cycles of PCR using NEBNext Ultra II Q5 master Mix (NEB #M0544L). Each PCR reaction contained a pool of forward primers (0.5 μM final concentration) and a unique indexed reverse primer (0.5 μM)(Bosch et al., 2021). Forward primers contain a P5 flow cell attachment sequence, a standard Read1 Illumina sequencing primer binding site, and custom stagger sequences to guarantee base diversity during Illumina sequencing. Reverse primers contain a P7 flow cell attachment sequence, a standard Read2 Illumina sequencing primer binding site, and unique barcodes to allow for sample pooling during deep sequencing.

Following PCR amplification, each ∼230 bp amplicon was purified using AMPure XP beads (Beckman– Coulter #A63882) using one-sided selection (1.2 x). Bead-purified amplicons were further purified on a Pippin HT 2% agarose gel cassette (target range 180-250 bp; Sage Science #HTC2010) to remove carry-over primer and genomic DNA. Eluted amplicons were quantified with Qubit 2.0 Fluorometer (Invitrogen) and amplicon size and purity was quality controlled by visualization on an Agilent 2100 Bioanalyzer (high sensitivity chip; Agilent Technologies #5067-4626). Next, individual PCR amplicons were multiplexed into 10 nM pools and sequenced on an Illumina sequencer according to the manufacturer’s instructions. To increase sequencing diversity, a PhiX spike-in of 2.5-5% was added to the pools (PhiX Sequencing Control v3; Illumina # FC-110-3001). Samples were run on the Illumina NextSeq 500, HiSeq 2500, or NovaSeq 6000 platform (Single-Read 1 x 85 cycles and 6 x i7 index cycles).

### NGS data processing, analysis, and hit calling

Sequencing counts were obtained in the manner described by(Bosch et al., 2021). Counts were normalized for sequencing depth and an sgRNA limit of detection (LOD) cut-off was set at 100 counts in the DMSO condition. Only sgRNAs that made the LOD cut-off (i.e. counts > 100) were analyzed further. Replicate screens were quality controlled to ensure that the Pearson correlation was >0.95 for both the non-targeting sgRNA sets and essential-gene targeting sgRNA(Dejesus et al., 2017) sets between each replicate screen. sgRNA counts were analyzed using MAGeCK analysis method (version 0.5.9.2) in python (version 2.7.16)(Li et al., 2014). Gene-level log2 fold change (L2FC) was calculated using the ‘alphamedian’ approach specified with the ‘gene-lfc-method’ parameter, which estimates the gene-level L2FC as the median of sgRNAs that are ranked above the default cut off in the Robust Rank Analysis used by MAGeCK. Negative control sgRNAs in the library were used to calculate the null distribution and to normalize counts using the ‘--control-sgrna’ and ‘–normalization control’ parameters, respectively. MAGeCK gene summary output results can be found in **Supplemental Data 1**.

Unless otherwise specified, a gene was determined to be a hit in a given condition if it had an FDR < 0.01 and a L2FC < −1 in the negative selection or an FDR < 0.01 and a L2FC > 1 in the positive selection.

### Clustered heatmap

L2FC values from the MAGeCK output (**Supplemental Data 1**) were used for the generation of clustered heatmap. Genes were clustered based on Euclidean distance using the Ward clustering criterion. Only genes that hit (FDR < 0.01; n=676 genes) in two or more conditions are included in the heatmap. A gene’s L2FC was only represented on the heatmap if the FDR was below 0.01 in the specific treatment condition; genes not meeting this significance threshold are shown as white in that treatment condition. Treatment conditions were clustered based on Pearson correlation using the Ward criterion. Clustering was done using the package hclust and the heatmap was generated using the heatmap.2 function from the package gplots (R version 4.0.5).

### Physiochemical property analysis

The physiochemical properties shown in **Supplemental Data 3D** were obtained from PubChem (https://pubchem.ncbi.nlm.nih.gov/) and, where applicable, were computed by Cactvs 3.4.8.18. The L2FC distributions correspond to the values for the following mAGP-associated genes found in (Jankute et al., 2015; Maitra et al., 2019): *rv1302, rv3265c, rv3782, rv3808c, rv1017c, rv3806c, rv3807c, rv3790, rv3791, rv3792, rv3794, rv3795, rv2673, rv0236c, rv3805c, rv3631, rv3779, rv2524c, rv3285, rv3280, rv0904c, rv2247, rv1483, rv2483, rv0533c, rv3799c, rv2245, rv2246, rv1483, rv0635, rv0636, rv0637, rv1484, rv0645c, rv0644c, rv0643c, rv0642c, rv0470c, rv3392c, rv0503c, rv3801c, rv3800c, rv2509, rv0206c, rv3804c, rv3802c, rv3436c, rv3441c, rv1018c, rv1315, rv0482, rv2152c, rv2155c, rv2158c, rv2157c, rv2156c, rv2153c, rv3818, rv3712, rv3713, rv2201, rv3910, rv2154c, rv0050, rv3682, rv3330, rv2911, rv0116c, rv2518c, rv1433, rv0192, rv0483, rv1477, rv1478, rv2190c, rv0867c, rv1009, rv1884c, rv2389c, rv2450c*.

### Antibacterial activity measurements

All compounds were dissolved in DMSO (VWR V0231) and dispensed using an HP D300e Digital Dispenser in a 384 well plate format. DMSO did not exceed 1% of the final culture volume and was maintained at the same concentration across all samples. CRISPRi strains were growth-synchronized and pre-depleted in the presence of ATc (100 ng/mL) for 5 days prior to assay for MIC analysis. Cultures were then back diluted to a starting OD_580_ of 0.05 and 50 µL of cell suspension was plated in technical triplicate in wells containing the test compound and fresh ATc (100 ng/mL). For checkerboard assays and MIC assays (non-CRISPRi) cultures were growth-synchronized to late log-phase and back-diluted to an OD_600_ of 0.025 prior to plating (no ATc). Plates were incubated at 37°C with 5% CO_2_. OD_600_ was evaluated using a Tecan Spark plate reader at 10-14 days post-plating and percent growth was calculated relative to the vehicle control for each strain. IC_50_ measurements were calculated using a non-linear fit in GraphPad Prism.

To quantify growth phenotypes on 7H10 agar, 10-fold serial dilutions of Mtb cultures (OD_600_ =0.6) were spotted on 7H10 agar containing drugs at the indicated concentrations and/or ATc at 100 ng/mL. Plates were incubated at 37°C and imaged after two weeks.

### Antimicrobial compounds

All compounds used in this study were purchased from commercial manufacturers with the exception of GSK3011724A, which was synthesized at the Memorial Sloan Kettering Organic Synthesis core as described in(Kumar et al., 2018).

| Antibiotic | Use | Abbreviation | Product number |
| --- | --- | --- | --- |
| Amikacin | Axenic culture | AMK | A0365900 (Sigma Aldrich) |
| Anhydrotetracycline | Axenic culture | ATc | AC233135000 (Fisher Scientific) |
| Azithromycin | Mouse efficacy studies | AZM | Epic Pharma Oral Suspension |
| Azithromycin-dihydrate | Axenic culture | AZM | PZ0007 (Sigma Aldrich) |
| Bedaquiline | Axenic culture | BDQ | AdooQ Biosciences (A12327-5) |
| Capreomycin Sulfate | Axenic culture | CAP | PHR1716 (Sigma Aldrich) |
| Chloramphenicol | Axenic culture | CHL | C-105-5 (GoldBio) |
| Clarithromycin | Axenic culture | CLR | C9742 (Sigma Aldrich) |
| Clarithromycin | Mouse efficacy studies | CLR | Sandoz Tablets |
| Clindamycin hydrochloride | Axenic culture | CLI | C-175-10 (GoldBio) |
| Clofazimine | Axenic culture | CLO | TCC2866 (VWR) |
| Ethambutol dihydrochloride | Axenic culture | EMB | E4630 (Sigma Aldrich) |
| Fusidic acid sodium salt | Axenic culture | FA | F0881 (Sigma Aldrich) |
| GSK3011724A | Axenic culture, macrophage infection | GSK'724A | NA |
| Isoniazid | Axenic culture | INH | I3377 (Sigma Aldrich) |
| Kanamycin | Axenic culture | KAN | K-120-50 (GoldBio) |
| Levofloxacin | Axenic culture | LFX | 28266 (Sigma Aldrich) |
| Lincomycin hydrochloride | Axenic culture | LNC | 62143 (Sigma Aldrich) |
| Linezolid | Axenic culture | LZD | SML1290 (Sigma Aldrich) |
| Moxifloxacin | Axenic culture | MXF | Y0000703 (Sigma Aldrich) |
| Rifampicin | Axenic culture, macrophage infection | RIF | R0079 (TCI) |
| Rifampicin | Mouse efficacy studies | RIF | Akorn Capsules |
| Streptomycin sulfate | Axenic culture | STR | S-150-100 (GoldBio) |
| Sutezolid | Axenic culture | SZD | PZ0035 (Sigma Aldrich) |
| Telithromycin | Axenic culture | TLM | SML2162 (Sigma Aldrich) |
| Tigecycline hydrate | Axenic culture | TGC | PZ0021 (Sigma Aldrich) |
| Vancomycin hydrochloride | Axenic culture | VAN | V2002 (Sigma Aldrich) |
| Zeocin | Axenic culture | ZEO | J67140-8EQ (Alfa Aesar) |

### Bone marrow-derived macrophage infections

Bone marrow-derived macrophages (BMDMs) were differentiated from wild-type, female C57BL/6NTAC mice (Taconic Farms, 6-8 weeks of age). All animal work was performed in accordance with the Guide for the Care and Use of Laboratory Animals of the National Institutes of Health, with approval from the Institutional Animal Care and Use Committee of Rockefeller University. Femurs and tibias were harvested and crushed with a sterile mortar and pestle as described(Trouplin et al., 2013). After red blood cell lysis and counter-selection of resident macrophages, bone marrow cells were incubated in the presence of DMEM (4.5 g/L glucose + L-glutamine + sodium pyruvate, Corning 10-013-CV) + 10% FBS (Sigma Aldrich F4135, Lot no. 17B189) + 15% conditioned L929 cell medium (LCM) and differentiated for 7 days at 37°C, 5% CO_2_. Macrophages were then lifted using gentle cell scraping. For infection assays, BMDMs were seeded in 96 well plates at 75,000 cells/well two days prior to infection. 16 hours prior to infection, fresh DMEM + 10% FBS + 10% LCM was added to cells, with or without IFN-γ (20 ng/mL, Gemini Biosciences 300-311P). Mtb cultures were synchronized to late log-phase (OD_600_ 0.6-0.8). For infections with CRISPRi strains, cultures were pre-depleted with 100 ng/mL ATc for 24 hours prior to infection. Mtb pellets were washed with PBS (Thermo Fisher 14190144) + 0.05% Tyloxapol (Sigma Aldrich T0307) and single cell suspensions were generated by harvesting the suspended cells after gentle centrifugation (150 x g for 12 min). Cell culture medium was removed from the macrophages and replaced with Mtb-containing medium at a multiplicity of infection of MOI of 1:1. After four hours of infection at 37°C, media was removed and cells were washed 2X with PBS. Wells were replenished with fresh media with or without drug. DMSO was normalized to 0.2%. For CRISPRi infections, doxycycline (Sigma Aldrich D9891) was added at a concentration of 250 ng/mL to maintain target knockdown. For all infection assays, medium was replaced with fresh drug at day 3. At each indicated timepoint, after two PBS washes, cells were lysed with 100 µL of Triton X-100 in water (Sigma Aldrich X100). Lysates were titrated in PBS + 0.05% Tween-80 and plated on 7H10. CFU were enumerated after 21-28 days of outgrowth.

### Selection of drug-resistant Mtb isolates

For the selection of linezolid-resistant H37Rv Mtb mutants, two independent cultures were started at an OD_600_ of 0.001. After one week of outgrowth, cultures were pelleted and roughly 3×10^9^ CFU were plated on complete 7H10 + 11.9 µM linezolid. Plates were incubated for 24 days. Colonies were picked and grown in complete 7H9 + 11.9 µM linezolid. Genomic DNA was harvested as described above? And Sanger sequencing was performed on purified PCR amplicons of *rrl* (23s rRNA) and *rplC* using the primers listed **(Supplemental Table 1)**. Genomic DNA was also submitted for WGS (BGI, llumina HiSeq X Ten platform).

For selection of resistant isolates for the lineage 1.2.1 strain, thirty independent cultures of 20 mL were started at an OD_600_ of 0.001. After growth to log-phase (OD_600_ 0.5-0.6) cultures were pelleted and plated on complete 7H10 + antibiotic (CLR = 10 µg/mL, RIF = 0.5 µg/mL). After 28-35 days, colonies were enumerated. CLR-resistant colonies occurred at a frequency of 1.7 x 10^-8^ and were picked and grown in complete 7H9 + CLR (4 µg/mL). Samples were heat lysed and *whiB7* and *rrl* were PCR amplified and sequenced using the primers listed **(Supplemental Table 1)**. Sanger sequencing revealed a wild-type *whiB7* locus in all (n=101) clarithromycin-resistant isolates. Instead, clarithromycin resistance was conferred by a variety of base substitutions in the 23S rRNA (**Supplemental Figure 10C,D**). Select samples were cultured further and purified genomic DNA was submitted for WGS.

### Total RNA extraction and qRT-PCR

Total RNA extraction was performed as previously described (Bosch et al., 2021). Briefly, 2 OD_600_ units of bacteria were added to an equivalent volume of GTC buffer (5M guanidinium thiocyanate, 0.5% sodium N-lauroylsarcosine, 25 mM trisodium citrate dihydrate, and 0.1M 2-mercaptoethanol), pelleted by centrifugation, resuspended in 1 mL TRIzol (Thermo Fisher Scientific; #15596026) and lysed by zirconium bead beating (MP Biomedicals; #116911050). 0.2 mL chloroform was added to each sample and samples were frozen at −80°C. After thawing, samples were centrifuged to separate phases, and the aqueous phase was purified by Direct-zol RNA miniprep (Zymo Research; # R2052). Residual genomic DNA was removed by TURBO DNase treatment (Invitrogen Ambion; # AM2238). After RNA cleanup and concentration (Zymo Research; #R1017), 3 µg of RNA per sample was reverse transcribed into cDNA with random hexamers (Thermo Fisher Scientific; # 18-091-050) following manufacturer’s instructions. RNA was removed by alkaline hydrolysis and cDNA was purified with PCR clean-up columns (Qiagen; #28115). Next, knockdown of the targets was quantified by SYBR green dye-based quantitative real-time PCR (Applied Biosystems; #4309155) on a Quantstudio System 5 (Thermofisher Scientific; #A28140) using gene-specific qPCR primers (5 µM), normalized to *sigA* (*rv2703*) and quantified by the ΔΔCt algorithm. All gene-specific qPCR primers were designed using the PrimerQuest tool from IDT (https://www.idtdna.com/PrimerQuest/Home/Index) and then validated for efficiency and linear range of amplification using standard qPCR approaches. Specificity was confirmed for each validated qPCR primer pair through melting curve analysis.

### RNA-seq cDNA library construction and deep sequencing

Triplicate cultures were grown to mid-log phase in 7H9 and diluted back to OD_600_=0.2 in 7H9 in the presence or the absence of ATc (100 ng/mL). Cultures were incubated for 48 hours, after which total RNA was extracted as described in “*Total RNA extraction and qRT-PCR*.” Following RNA cleanup (Zymo Research; #R1017), 2 μg total RNA for each sample was depleted for rRNA using a Ribominus Transcriptome Isolation Kit (Yeast and Bacteria, Invitrogen, K1550-03). Following rRNA depletion, RNA was concentrated using an RNA Clean and Concentration-5 kit (Zymo Research, R1013). RNA quality was then confirmed by Bioanlayzer (Agilent RNA 6000 Pico kit, 5067-1513).

We used the NEB Next Ultra II Directional RNA Library Prep Kit (NEB, E7760 and E7765) to prepare cDNA libraries, following manufacturer’s instructions. Briefly, 150 ng of rRNA-depleted RNA was subjected to fragmentation by incubating samples at 94°C for 20 min, followed by first strand cDNA synthesis (10 minutes at 25°C, 50 minutes at 42°C, 15 minutes at 70°C, hold at 4°C). Second-strand synthesis was performed at 16°C for 1.5 hours. DNA purification was performed with AMPure XP beads (Beckman Coulter, A63881). End repair was performed for 30 minutes at 20°C, followed by 30 minutes at 65°C. Repaired dsDNA was adaptor ligated (15 minutes at 37°C) and purified with AMPure XP beads. Eluted DNA was amplified by PCR using NGS primers supplied with the kit (NEBNext Multiplex Oligos for Illumina, Index Primers Set 1 and 2, E7335S, E7500S) for 12 cycles of amplification. Amplicons were purified with AMPure XP beads, quantified by Qubit dsDNA HS Assay kit (TheromoFisher Scientific, Q32851), and quality controlled by BioAnalzer (Agilent DNA 1000, 5067-1504). RNAseq libraries were sequenced on an Illumina NextSeq 500 (mid-output, 75 bp paired-end read).

### Processing and analysis of RNA-seq data

Raw FASTQ files were aligned to the H37Rv genome (NC_018143.2) using Rsubread (version 2.0.1)(Liao et al., 2019) with default settings. Transcript abundances were calculated by processing the resulting BAM files with the summarizeOverlaps function of the R package GenomicAlignments (version 1.22.1)(Lawrence et al., 2013). Overlaps were calculated in the “Union” mode, ensuring reads were counted only if they overlap a portion of a single gene/feature. 16S, 23S, and 5S rRNA features (RVBD6018, 6019, and 6020, respectively) were manually removed from the count data to prevent confounding downstream differential gene expression analysis. Differential expression analysis was conducting using DESeq2 (version 1.30.1)(Love et al., 2014) with default parameters.

### Cell wall permeability assay

Cell envelope permeability was determined using the ethidium bromide (EtBr) uptake assay as previously described(Xu et al., 2017). Briefly, mid-log-phase Mtb cultures were washed once in PBS + 0.05% Tween-80 and adjusted to OD_600_=0.8 in PBS supplemented with 0.4% glucose. 100 µL of bacterial suspension was added to a black 96-well clear-bottomed plate (Costar). After this, 100 μL of 2 μg/mL EtBr in PBS supplemented with 0.4% glucose was added to each well. EtBr fluorescence was measured (excitation: 530 nm/emission: 590 nm) at 1 min intervals over a course of 60 min. Experiments were performed in technical triplicate.

A similar assay was performed to determine envelope permeability to a fluorescent vancomycin analogue, except that: (1) the bacterial suspension was adjusted at OD_600_=0.4 in PBS supplemented with 0.4% glucose; (2) cells were incubated with 2 μg/mL BODIPY FL Vancomycin (Thermo Scientific, V34850); (3) 200 μL sample aliquots were taken at different time points, washed twice with PBS, resuspended in 200 μL PBS; and (4) fluorescence was measured (excitation: 485 nm/emission: 538 nm) and normalized to the OD_600_ of the final bacterial suspension.

### Whole genome sequencing data aggregation, alignment, SNP calling and annotation

FASTQ data were downloaded from NCBI using the SRA Toolkit (version 2.9.6). A list of accession numbers of all analyzed FASTQ files is provided in **Supplemental Data 4**. FASTQ reads were aligned to the H37Rv genome (NC_018143.2) and SNPs were called and annotated using Snippy (version 3.2-dev) using default parameters (minimum mapping quality of 60 in BWA, SAMtools base quality threshold of 20, minimum coverage of 10, minimum proportion of reads that differ from reference of 0.9 (Seemann, 2020). Mapping quality and coverage was further assessed using QualiMap with the default parameters (version 2.2.2-dev)(Okonechnikov et al., 2016). Samples with a mean coverage < 30, mean mapping quality <= 45, or GC content <= 50% or >= 70% were excluded. Spoligotypes were assigned using SpoTyping (version 2.1)(Xia et al., 2016). Drug resistance conferring SNPs were annotated using reference SNP lists from (Allix-Béguec et al., 2018; Sandgren et al., 2009) and Mykrobe v0.9.0 (Hunt et al., 2019).

Phylogenetic trees were built using FastTree (version 2.1.11 SSE3) (Price et al., 2010). A list of SNPs in essential genes was concatenated for the building phylogenetic trees. Indels, drug resistance-conferring SNPs, and SNPs in repetitive regions of the genome (PE/PPE genes, transposases and prophage genes) were excluded. Tree visualization was performed in iTol (https://itol.embl.de/).

### Sublineage identification

Mtb sublineages were assigned to each sample using a set of lineage identifying SNPs. Lineage identifying SNPs from(Coll et al., 2014; Palittapongarnpim et al., 2018) were combined and then reduced to a subset of synonymous SNPs occurring in essential genes. For each sample, the percentage of lineage identifying SNPs present was calculated for each possible sublineage. A threshold of 67% was set as the minimum percentage of sublineage identifying SNPs required in order to define a sublineage. For each sample, all sublineages meeting this threshold were then evaluated to determine if they formed a continuous line of descent from the highest sublineage to the lowest (e.g. lineage1 → lineage1.2 → lineage1.2.1 → lineage1.2.1.1). Samples with a continuous line of descent were assigned the most specific sublineage (e.g. lineage1.2.1.1). If the set of sublineages included other sublineages that did not fit within the line of descent, the sublineage call was marked as “not confident” and considered as an undetermined sublineage.

### Mouse infection and drug treatment

Female BALB/c mice (Charles Rivers Laboratory) 7-8 weeks old were infected with 100-200 CFU of Mtb using a whole-body inhalation exposure system (Glas-Col). After 10 days (**Figure 6D, E)** or 14 days **(Supplemental Figure 10H, I)**, animals were randomly assigned to study groups and chemotherapy was initiated. CLR and RIF were stirred in 0.5% CMC/0.5% Tween-80 to resuspend; INH was resuspended in water. AZM was resuspended in water. Liquid drug formulations were administered once daily by oral gavage for 14 consecutive days. After 13 days of drug treatment, blood samples of the mice were taken 1 hour and 24 hours post dosing (**Supplemental Figure 10J,K**). At designated time points (14 after starting chemotherapy), mice were euthanized, and lungs and spleens were aseptically removed, homogenized in 1 mL PBS + 0.05% Tween-80 and plated on Middlebrook 7H11 agar supplemented with 10% OADC. Colonies were counted after 4-6 weeks of incubation at 37°C. Mice were housed in groups of 5 in individually ventilated cages inside a certified ABSL-3 facility and had access to water and food *ad libitum* for the duration of the study. All experiments involving animals were approved by the Institutional Animal Care and Use Committee of the Center for Discovery and Innovation.

### Drug quantitation in plasma by high pressure liquid chromatography coupled to tandem mass spectrometry (LC-MS/MS)

Neat 1 mg/mL DMSO stocks for rifampicin (RIF), azithromycin (AZM), and clarithromycin (CLR) were serial diluted in 50/50 (acetonitrile/water) to create neat spiking stocks. Standards and quality controls were created by adding 10 µL of spiking stock to 90 µL of drug free plasma. 10 µL of control, standard, quality control, or study sample were added to 100 µL of 50/50 (acetonitrile/methanol) protein precipitation solvent containing the stable labeled internal standards RIF-d8 (Toronto Research Chemicals; R508003), AZM-d5 (Toronto Research Chemicals; A927004) and CLR-13C-d3 (Cayman Chemical; 26678) at 10 ng/mL. Extracts were vortexed for 5 min and centrifuged at 4,000 rpm for 5 min. 100 µL of supernatant of RIF containing samples was combined with 5 µL of 75 mg/mL ascorbic acid to stabilize RIF. 100 µL of mixture was combined with 100 µL of Milli-Q water prior to HPLC-MS/MS analysis. CD-1 mouse control plasma (K_2_EDTA) was sourced from Bioreclamation. RIF, AZM, and CLR were sourced from Sigma Aldrich.

LC-MS/MS analysis was performed on a Sciex Applied Biosystems Qtrap 6500+ triple-quadrupole mass spectrometer coupled to a Shimadzu Nexera X2 UHPLC system to quantify each drug in plasma. Chromatography was performed on an Agilent SB-C8 (2.1 x 30 mm; particle size, 3.5 µm) using a reverse phase gradient. Milli-Q deionized water with 0.1% formic acid was used for the aqueous mobile phase and 0.1% formic acid in acetonitrile for the organic mobile phase. Multiple-reaction monitoring of precursor/product transitions in electrospray positive-ionization mode was used to quantify the analytes. Sample analysis was accepted if the concentrations of the quality control samples were within 20% of the nominal concentration. The compounds were ionized using ESI positive mode ionization and monitored using masses RIF (823.50/791.60), AZM (749.38/591.30), CLR (748.38/158.20), RIF-d8 (831.50/799.60), AZM-d5 (754.37/596.30), and CLR-d4 (752.33/162.10). Data processing was performed using Analyst software (version 1.6.2; Applied Biosystems Sciex).

## DATA AVAILABILITY

Raw sequencing data will be deposited to the Short Read Archive (SRA) under project number PRJNA738381. All screen results are available in **Supplemental Data 1** and at pebble.rockefeller.edu.

